# The *autophagy-related genes AtATG5* and *AtATG7* influence reserve mobilisation and response to ABA during seed germination

**DOI:** 10.1101/2024.05.15.593177

**Authors:** Estefanía Contreras, Inmaculada Sánchez-Vicente, Elena Pastor-Mora, Mar Aylón-Rodríguez, Mar G. Ceballos, Miguel Ángel Delgado-Gutiérrez, Óscar Lorenzo, Jesús Vicente-Carbajosa, Raquel Iglesias-Fernández

## Abstract

Autophagy is an intracellular recycling mechanism that generally degrades cytoplasmic components non-selectively, but it can also target specific substrates under certain conditions. Here, we investigate the impact of autophagy on Arabidopsis seed biology through autophagy-related (*ATG*) genes *AtATG5* and *AtATG7*, and their role in ABA responses. Seeds of *atg5* and *atg7* mutants germinate significantly slower than Col-0, especially under ABA, and show transcriptomic differences and histochemical alterations in the organization of lipid droplets and protein storage vacuoles. Notably, immunolocalization of ATG8 is observed in PSV of Col-0, but not in *atg* mutants. Differentially expressed genes in *atg7* compared to Col-0 in response to ABA include reported targets of the transcription factor (TF) ABI5, a master regulator of ABA signalling in the seed. Interestingly, the decrease in ABI5 normally observed in Col-0 seeds after imbibition is delayed in *atg* mutants, which also show altered accumulation in developing seeds of the bZIP67, an ABI5 homolog that regulates reserve biosynthesis. Yeast two-hybrid and co-immunoprecipitation assays confirmed a direct interaction with the autophagy machinery *in vitro* and *in vivo*, mediated through ATG8. Our data highlight the relevance of autophagy in seed reserve mobilization, and seed germination through ABA responses by a TF decay mechanism.

## Introduction

Autophagy is a conserved recycling mechanism in eukaryotes that degrades cytoplasmic components, removing unnecessary or damaged elements. While traditionally seen as a non-selective process, it also targets specific proteins, lipids, and organelles in response to biotic and abiotic stresses (Su *et al*., 2020; Qi *et al*., 2021). Plant autophagy plays a crucial role in cellular homeostasis and facilitates adaptation to changing environments, making it essential for plant survival under various stresses and developmental programs (Liu *et al*., 2009; Di Berardino *et al*., 2018). In plant cells, autophagy occurs in the central vacuole via vesicular trafficking, mainly through macroautophagy and microautophagy. The most studied, macroautophagy, transports cargo in double-membrane autophagosomes from the endoplasmic reticulum (ER) to the vacuole (Michaeli *et al*., 2016). In microautophagy, cytoplasmic elements such as lipid droplets (LD; also known as oil bodies and oleosomes), specialized organelles accumulating triacylglycerols (TAG) are directly engulfed by the central vacuole through tonoplast invagination (Siénko *et al*., 2020).

In Arabidopsis, autophagy involves around 40 conserved autophagy-related proteins (ATGs) and other evolutionary conserved factors from over 17 protein families, including kinases, ubiquitin-like proteins, vacuolar sorting factors, and cysteine proteases (Ding *et al*., 2018). *In A. thaliana*, ATGs are involved in three main complexes for autophagosome formation: (i) the ATG1 kinase complex, which induces autophagy via interaction with the Target Of Rapamycin (TOR) under nutrient starvation (Dobrenel *et al*., 2016); (ii) the phosphatidylinositol 3-kinase (PI3K) complex, which initiates phagophore nucleation (Yang *et al*., 2010); and (iii) the ATG9–ATG18–ATG2 complex, which recruits membranes for phagophore expansion (Sawa-Makarska *et al*., 2020). The ATG8– ATG12 conjugation system attaches phosphatidylethanolamine (PE) to ATG8, enabling selective cargo recognition via ATG8-interacting motifs (AIMs) that bind to selective receptors like Neighbour of BRCA1 gene 1 (NBR1) and translocator protein (TSPO) (Kellner *et al*., 2017; Ding *et al*., 2018; Luo *et al*., 2021). The autophagy-related factors AtATG5 and AtATG7 are crucial for the conjugation of PE to ATG8 in both animals and plants (Xie and Klionsky, 2007; Minina *et al*., 2018; Fraiberg and Elazar, 2020). Genetic studies in *A. thaliana* have shown that *ATG5* and *ATG7* are essential for storage protein and lipid accumulation during embryogenesis and seed maturation. Their overexpression enhances seed production and nitrogen content, while *atg5* and *atg7* mutants display altered seed composition (Guiboileau *et al*., 2013; Masclaux-Daubresse *et al*., 2014; Di Berardino *et al*., 2018; Minina *et al*., 2018; Fan *et al*., 2019). Similarly, in *Oryza sativa*, autophagy affects starch quality, influencing both seed size and appearance, as *Osatg7-1* produces fewer and small seeds with a chalky appearance and a low starch content in endosperm (Sera *et al*., 2019; Iglesias-Fernández & Vicente-Carbajosa, 2022).

Seeds are essential for plant evolution, enabling dispersal, survival under harsh conditions, and regrowth in favourable environments (Finch-Savage & Bassel, 2016). Seed maturation, germination, and post-germination involve the accumulation of reserves like starch, seed storage proteins (SSP), and lipids (triacylglycerols; TAG) (Holdsworth *et al*., 2008; Bewley *et al*., 2013; Sajeev *et al*., 2024). SSP synthesis and storage in maturing seeds occur in the ER, which transports them to seed-specific vacuoles, known as Protein Storage Vacuoles (PSV), through an autophagy-like process (Hills, 2004; Vitale *et al*., 2022). Seed germination starts with water uptake and ends with the breakage of embryo-covering layers (i.e. endosperm rupture) that allows radicle emergence (germination *sensu stricto*). The post-germination stages involve the hydrolysis of seed storage reserves by *de novo* synthesised proteases, providing essential support for seedling growth until the photosynthetic capacity is acquired (Nakabayashi *et al*., 2005). In imbibed Arabidopsis seeds, the degradation of SSP begins at the radicle, coinciding with cell expansion and differentiation. After radicle emergence (i.e. post-germination period), SSP catabolism occurs mainly in the cotyledons, especially when the SSP reserves of the embryonic axis are nearly depleted (Müntz, 2007; Iglesias-Fernández *et al*., 2014). In *Vigna mungo* (Fabaceae), the ERvt (ER to vacuole trafficking) pathway has been reported to transport cysteine proteases from the ER to PSV, encapsulated within KDEL Vesicles (KV) (Okamoto *et al*., 2003). Interestingly, the autophagy receptor TSPO plays a major role in the regulation of fatty acids (FA) and LD in Arabidopsis germinating seeds and in seedlings. Recently, studies in the knock-out mutant of LD structural protein -CALEOSIN 1 (CLO1) have described its influence in the degradation of LD by autophagy (lipophagy) in germinating seeds of Arabidopsis (Jurkiewicz *et al*., 2018; Miklaszewska *et al*., 2023; Qin *et al*., 2023; Zhuang *et al*., 2024).

In natural soil seedbeds, seeds encounter a complex environment with various stresses, requiring mechanisms for survival and adaptation. Seed vigor, defined by the ability to germinate under different conditions, is crucial for this adaptation (Finch-Savage and Bassel, 2016). The ER plays a central role in detecting and managing stress responses. When stress occurs, the Unfolded Protein Response (UPR) pathway is activated to prevent protein accumulation in the ER and restore homeostasis (Ellgaard *et al*., 2003; Yang *et al*., 2016). Persistent stress leads to the selective removal of ER domains via autophagy, helping manage protein overload (Stephani *et al*., 2020). While autophagy is essential for degradation in response to various stimuli, its role in seeds remains underexplored. In *Medicago truncatula*, ER stress and autophagy, mediated by MtATGs, are key for seed development and drought response (Yang *et al*., 2021). Abscisic acid (ABA) regulates plant adaptation to abiotic stresses and controls seed maturation and dormancy. Environmental stress during embryo development can halt germination through an ABA-dependent mechanism. In *Arabidopsis thaliana*, the autophagy receptor NBR1 interacts with ABA signaling transcription factors (TFs: ABI3, ABI4, ABI5), and its overexpression reduces seed sensitivity to ABA (Carbonero *et al*., 2017; Tarnowski *et al*., 2020; Luo *et al*., 2021; Chen *et al*., 2022; Matilla, 2022).

This study investigates the role of autophagy in seed biology by analyzing *Arabidopsis thaliana AtATG5* and *AtATG7* loss-of-function mutants. Seeds from *atg5* and *atg7* knockout mutants germinate slower than wild type (WT, Col-0), both under normal conditions and with ABA. Histochemical analysis showed alterations in storage compounds (proteins and lipids) in these mutants compared to WT seeds at both dry and imbibed stages. Immunofluorescence revealed that ATG8a did not localize to PSV in autophagy-deficient mutants, supporting the role of autophagy in reserve mobilization. Transcriptomic analysis during germination identified significant transcriptional differences between *atg7* and WT seeds, particularly in ABA-imbibed seeds, with 22.5% of differentially expressed genes linked to ABI5 TF response. Western blot analysis showed altered levels of ABI5 and bZIP67 in autophagy mutants, indicating autophagy’s regulatory role in seed development and imbibition. Protein-protein interactions between ATG8 and ABI3/ABI5 were confirmed through yeast two-hybrid assays (Y2H). Co-IP assays have *in vivo* confirmed the ABI5-ATG8 interaction in Arabidopsis 4 day-old seedlings. Overall, the study highlights autophagy’s involvement in seed development, germination, and the seed’s response to environmental signals, such as ABA.

## Materials and Methods

### Plant material and growth conditions

*Arabidopsis thaliana* Columbia (Col-0) ecotype was used as WT. Knockout T-DNA insertion lines were used for *ATG5* (*atg5-1;* SAIL_129B079) and *ATG7* (*atg7-2;* GK-655B06; Hofius *et al*., 2009; Yoshimoto *et al*., 2009), kindly donated by C. Masclaux-Daubresse (IJPB, France), *ABI5* (*abi5-7*; Nambara *et al*., 2002; Albertos *et al*., 2015), *35S:cMyc-ABI5* (Albertos et al, 2015) and *ProUBQ10-GFP-ATG8A* (kindly provided by Dr. Cecilia Gotor and Dr. Luis C. Romero, Instituto de Bioquímica Vegetal y Fotosíntesis-CSIC-Sevilla; Calero-Muñoz *et al*., 2019; Nakamura *et al*., 2021).

Seeds were surface sterilized in 70% ethanol for 15 minutes, followed by a 100% ethanol wash and air-dried. They were then plated on solid half-strength Murashige and Skoog medium with 1% sucrose (MS/2, Duchefa Biochemie, Haarlem, Netherlands) and stratified at 4°C for 3 days. Seeds were incubated for 2 weeks at 21°C in a germination chamber under a long-day photoperiod (16h light/8h dark) with a light intensity of 155 μmol photons/m²/s. Seedlings were transferred to soil:vermiculite (3:1) pots and grown in a greenhouse under similar conditions. Siliques were collected at different seed embryogenesis and maturation stages for RNA isolation. Mature brown seeds were harvested, stored at 21°C and 30% relative humidity, and considered after-ripened after one month.

### Germination assays

Three replicates of 50 after-ripened, non-stratified mature seeds were sterilized and then imbibed on moist grid-patterned nitrocellulose filter paper (Whatman n°1; Thermo Fisher Scientific; Waltham, MA, USA) placed on 55 mm Petri dishes containing solid MS/2, with or without 1 μM 2-cis, 4-trans-(±) Abscisic acid (ABA; Merck, Rahway, NY, USA). Plates were kept in a germination chamber under the specified conditions. Seed Coat Rupture (SCR) and Endosperm Rupture were evaluated at 0, 24, 30, 36, 48, 54, 72, 76, and 80 hours of imbibition (hoi) using an Olympus stereo microscope SZ2-ILST (Olympus; Tokyo, Japan). Statistical analysis and the time to reach 50% germination (t50) were calculated using the GERMINATOR Microsoft Office Excel 16.9 package (Joosen et al., 2010). Col-0 and *atg7* imbibed seeds were also collected at selected time points (0, 24, 48, 96 hoi) in the absence and presence of 1 μM ABA and frozen in liquid nitrogen for RNA isolation and subsequent RNA-seq analysis.

### Generation of *PAtATG5::uidA* and *PAtATG7::uidA* transgenic lines and histochemical GUS assays

*AtATG5* (*At5g17290*) and *AtATG7* (*At5g45900*) native promoter sequences were amplified from *A. thaliana* genomic DNA (from -404 bp and -340 bp before the initial ATG, respectively) using oligonucleotides with *Bam*HI and *Sal*I restriction sites. PCR products were cloned into the pGEM T-easy vector (Thermo Fisher Scientific) and transferred to the pBI101 vector (between *Bam*HI and *Sal*I sites), which contains the *Escherichia coli uidA* reporter gene. Recombinant plasmids were transformed into C51C1 *Agrobacterium tumefaciens* cells via electroporation (ECM 630 Electroporator; BTX; Harvard Bioscience, Holliston, MA, USA) and introduced into *A. thaliana* Col-0 plants using Agrobacterium floral dip transformation (Clough and Bent, 1998). The presence of the pBI101 vector in T1 plants was confirmed by hygromycin resistance (40 mg/L). Plants from subsequent generations were self-fertilized until homozygous lines were identified in T3, with 100% germination on selection media. *AtATG5* and *AtATG7* promoter expression was evaluated in a representative transgenic line by qualitative GUS staining (Jefferson et al., 1987) and visualized under a Leica DMi8 light microscope (Leica; Wetzlar, Germany).

### RNA isolation, cDNA synthesis, and quantitative PCR assays (qPCR)

Total RNA was purified from Col-0, *atg5*, and *atg7* at different germination time points (0, 12, 24, 30, 36, 42 hours of imbibition) and seed stages (early green, late green, and brown seeds) as described by Oñate-Sánchez and Vicente-Carbajosa (2008). RNA quality and concentration were assessed using a Nanodrop® ND-1000 spectrophotometer (Thermo Fisher Scientific). cDNA was synthesized from 1 µg of RNA using the RevertAid First Strand cDNA Synthesis Kit (Thermo Fisher Scientific) following the manufacturer’s instructions, and samples were stored at -20°C. qPCR was performed on an Eco® Real-Time PCR System (Illumina, San Diego, CA, USA) using primers for *AtATG5*, *AtATG7*, and *AtACT8* (Graeber et al., 2011; Iglesias-Fernández et al., 2014) as a reference gene. The 10 µl reaction contained cDNA, FastStart SYBR Green Master (ROX; Hoffmann-La Roche, Basel, Switzerland), primers, and water. Thermal cycling conditions were 95°C for 10 min, followed by 40 cycles of 95°C for 10 sec and 60°C for 30 sec. A melting curve was used to calculate melting temperatures. Primer efficiencies (E) were estimated from cDNA serial dilutions, using the formula E[=[(10^−1^/regression line slope − 1)[×[100. Expression levels were determined by the cycle threshold (CT) in the exponential phase of PCR (Pfaffl, 2001).

### RNA library construction and RNA-seq analysis

Total RNA was isolated from Col-0 and *atg7* seeds, with RNA integrity confirmed using a Bioanalyzer 2100 (Agilent). Library construction and RNA sequencing (PE150) were performed on Illumina NovaSeq™ 6000 platforms (Illumina, San Diego, CA, USA) by Novogene Genomics Service (Novogene, Beijing, China) with three biological replicates per genotype and treatment. mRNA was purified from total RNA using poly-T oligo-attached magnetic beads, fragmented, and synthesized into cDNA. The cDNA was subjected to end repair, A-tailing, adapter ligation, amplification, and purification. Library quantification was done using Qubit and real-time PCR, with size distribution checked using a bioanalyzer. Clean reads were obtained by removing low-quality reads and adapters and aligned to *the A. thaliana* reference genome (TAIR10) using Hisat2 v2.0.5 (Kim *et al*., 2019; Anders *et al*., 2015). Differential expression analysis was conducted with DESeq2 (1.42.0; Love *et al*., 2014), with adjusted p-values (padj) ≤ 0.05 indicating differentially expressed genes. Principal component analysis (PCA) was performed using R (version 4.3.1), and visualization was done using the FactoExtra package. Gene set enrichment analysis (GSEA) was carried out with the gseGO function in the clusterProfiler package, with visualization via ridge plots using ggridges. Heatmaps were constructed using R studio and InstantClue v.0.10.10.dev-snap (Nolte *et al*., 2018) and Gene Ontology (GO) term enrichment analysis was done using ShinyGO 0.80 (Ge *et al*., 2020).

### Fixation, embedding, and sectioning of material for histochemistry

Arabidopsis Col-0, dry and imbibed (24, 48 hoi) seeds were infiltrated with formaldehyde: acetic acid: ethanol: water (3.5:5:50:41.5 by vol.) for 45 min under vacuum (41 mbar) and then incubated at 4°C for 3 days with gentle shaking as described by Ferrándiz *et al*. (2000) with modifications. Increasing concentrations of ethanol solutions were used to dehydrate sections and subsequently ethanol was progressively replaced with HistoClear (National Diagnostics, Hessle Hull, England) and then embedded in paraffin. Thin sections of 8 µm were performed with a Leica HistoCore NANOCUT R microtome (Leica) and collected on glass slides. Before histological staining and fluorescent immunolocalization, samples were de-waxed, by repeating the washes in reverse order.

### Fluorescent immunolocalization

Seed histological sections were washed with Phosphate Buffered Saline (PBS) and digested with 1 mg/ml proteinase-K (Hoffmann-La Roche). Slides were then incubated for 30 minutes at room temperature with a blocking solution (Blocking Reagent Roche), followed by overnight incubation with primary Anti-APG8A/ATG8A antibody (ab77003; Abcam, Cambridge, UK) at a 1:1000 dilution in 3% BSA and 0.5% Triton X-100 (Merck). After washing with Tris-Buffered Saline (TBS) containing 5 mM Na-Azide, sections were incubated for 3 hours with secondary Horse Anti-Rabbit IgG Antibody, DyLight™ 488 (DI-1088-1.5; Thermo Fisher Scientific) at a 1:100 dilution in the same buffer. Finally, samples were washed in PBS and water, mounted in CitifluorTM (Sciences Services GmbH, München, Germany), and examined using a Zeiss LSM 880 confocal microscope (excitation 493 nm, emission 512 nm).

### Polysaccharide and protein histological staining

Seed sections were stained with PAS-NBB: 0.5% (w/v) Periodic Acid (Merck) plus Schiff’s reagent (Merck) to detect polysaccharides (pink staining), and with 1% (w/v) Naphthol Blue Black (Merck) to visualize proteins (blue staining). Microscopy analyses were performed on a Zeiss LSM 880 microscope in bright field mode, and the images were captured and processed with the Zen Blue Edition software (Zeiss).

### Embryo lipid staining

The coat of Col-0, *atg5*, and *atg7* imbibed seeds (24, 48 hoi) was dissected using hypodermic needles, and embryos were collected in PBS. Then, they were fixed in 4% paraformaldehyde in PBS for 1h under vacuum (41 mbar), washed twice in PBS, and incubated o/n in ClearSee solution (10% xylitol, 15% sodium deoxycholate, 25% urea in distilled water) with gentle agitation. Embryos were stained o/n with 0.05% Nile Red in ClearSee, washed for 30 min and then 1h in ClearSee solution, and mounted in the same solution for imaging on a Zeiss LSM 880 microscope (excitation 561, emission 620 nm) and subsequently processed with Airyscan. Lipid quantification in embryo cells was analyzed using Fiji software (Schindelin *et al*., 2022) and has been expressed as a percentage of lipid area/cell area, in a minimum of 10 distinct embryos per line and germination time.

### Western blot

Total protein was extracted from 20 mg of Col-0, *atg5*, and *atg7* seeds at different germination time points (0, 36, 48, 72 hoi). Seeds were frozen in liquid nitrogen, ground using a Mikro-Dismembrator S (Sartorius AG, Göttingen, Germany), and mixed with 200 μL of extraction buffer (100 mM Tris-HCl pH 6.8, 4% SDS, 20% glycerol, 200 mM DTT). The samples were vortexed, heated at 95°C for 5 minutes, and centrifuged. Supernatants were recovered, and protein concentration was measured with the Bio-Rad Protein Assay (Bio-Rad, Hercules, CA, USA). Twenty-five μg of protein were analyzed using SDS-PAGE with a Mini-PROTEAN Tetra system (Bio-Rad), followed by transfer to a nitrocellulose membrane (Bio-Rad). Membranes were blocked in TBST with 5% BSA and probed with anti-ABI5 antibody (1:4,000; Agrisera, Vännäs, Sweden) overnight at 4°C. After washing, membranes were incubated with goat anti-rabbit ALP-conjugated antibody (1:10,000; Agrisera) for 1 hour. Detection was done using NBT/BCIP solution (Merck), and band intensity was quantified using ImageJ software.

For western blot analysis of siliques, total protein was extracted from green siliques of entire stems as described in Sánchez-Vicente *et al*., 2024. The tissue was homogenized in extraction buffer (100mM Tris-HCl, 150mM NaCl, 0.25% NP-40) with 1mM PMSF and 1X protease inhibitor. After centrifugation, protein concentration was quantified, and 70 μg were loaded on an SDS-PAGE. Proteins were transferred to a PVDF membrane (Millipore) and probed with antibodies, including anti-bZIP67 (Biomedal), anti-Actin (Sigma-Merck), and ECL-Peroxidase-labelled anti-rabbit and anti-mouse antibodies (Amersham). Detection was done using the ECL Advance kit (Amersham) and chemiluminescence was detected with the ChemiDoc Imaging System (Bio-Rad).

### Yeast two-hybrid (Y2H) assays

For Y2H assays, a mating-based yeast method and the *HIS3* reporter gene were used. A *Saccharomyces cerevisiae* J694α (α mating type) library containing a plasmid (pDEST™22, GAL4 activation domain, AD; Thermo Fisher Scientific) with 1,200 Open Reading Frames of TFs in *A. thaliana* was employed (Castrillo et al., 2011). Yeasts containing AtABI3 and AtABI5 were selected as prey. The coding sequence of *AtATG8* was amplified from *A. thaliana* cDNA, subcloned into pENTR3C, and then cloned into the pDEST™32 vector (GAL4 binding domain, BD) by Gateway strategy. This construct was used to transform *S. cerevisiae* YM4271a (a mating type) by PEG method and used as bait. Diploid cells were grown on solid DOB media (Bioworld, Dublin, OH, USA) lacking leucine and tryptophan. If AtATG8 interacts with AtABI3 and AtABI5 fused to BD, GAL4 TF is reconstituted, and Histidine (H) auxotrophy occurs. Increasing concentrations of 3-amino-1,2,4-triazole (3-AT; Merck) were added to avoid leakage of the *HIS*3 gene. Positive colonies appeared after 2-5 days of incubation at 28°C.

### Semi *in vivo* co-immunoprecipitation (CoIP) and pull-down assays

For CoIP assays, proteins were extracted from transgenic lines *35S:cMyc-ABI5* and *ProUBQ10-GFP-ATG8A.* Seedlings were grown in MS and then treated or not with 3-MA 5mM for 24 hours. Proteins were extracted from 4-day-old Arabidopsis seedlings with a buffer containing 50mM Tris– HCl, pH 7.5, 250mM NaCl, NP40 1%, 1mM PMSF and 1X cOmplete® EDTA-free proteases inhibitors (Sigma). Extracts were cleared by centrifugation and protein concentration was determined by Bradford assay. 1,5 mg of total soluble proteins were maintained in darkness at room temperature for 30 min. After treatment, extracts were incubated for 30 min in ice and then GFP-tag was magnetically labeled and immunoprecipitated using µMACS GFP Isolation Kit (MiltenyiBiotec) following manufacturer’s protocol. The proteins were visualized using anti-ABI5 (Biomedal, 1:10,000), anti-GFP-HRP (MiltenyiBiotec, 1:1,000) and ECL-Peroxidase-labelled anti-rabbit (Amersham, 1:10,000) antibodies.

## Results

### The *AtATG5* and *AtATG7* genes are expressed in embryos of *Arabidopsis thaliana* during seed maturation and germination

To study the impact of autophagy on seed biology, a selection of 68 *A. thaliana* autophagy genes was analyzed based on GO classification (see Materials and Methods). Their expression patterns were obtained from the Bio-Analytic Resource for Plant Biology (BAR) public database, and a heatmap representation was created (Fig. S1; Data S1). Among these genes, *ATG5* and *ATG7* were chosen for further study, as they are single-copy genes in Arabidopsis (*AtATG5*: *At5g17290*, *AtATG7*: *At5g45900*), and are essential components of the autophagy machinery in both plants and animals (Xie and Klionsky, 2007; Minina *et al*., 2018; Fraiberg and Elazar, 2020). Both genes showed high expression during late silique maturation and dry seed stages (DS) (Fig. 1a).

**Fig. 1.**
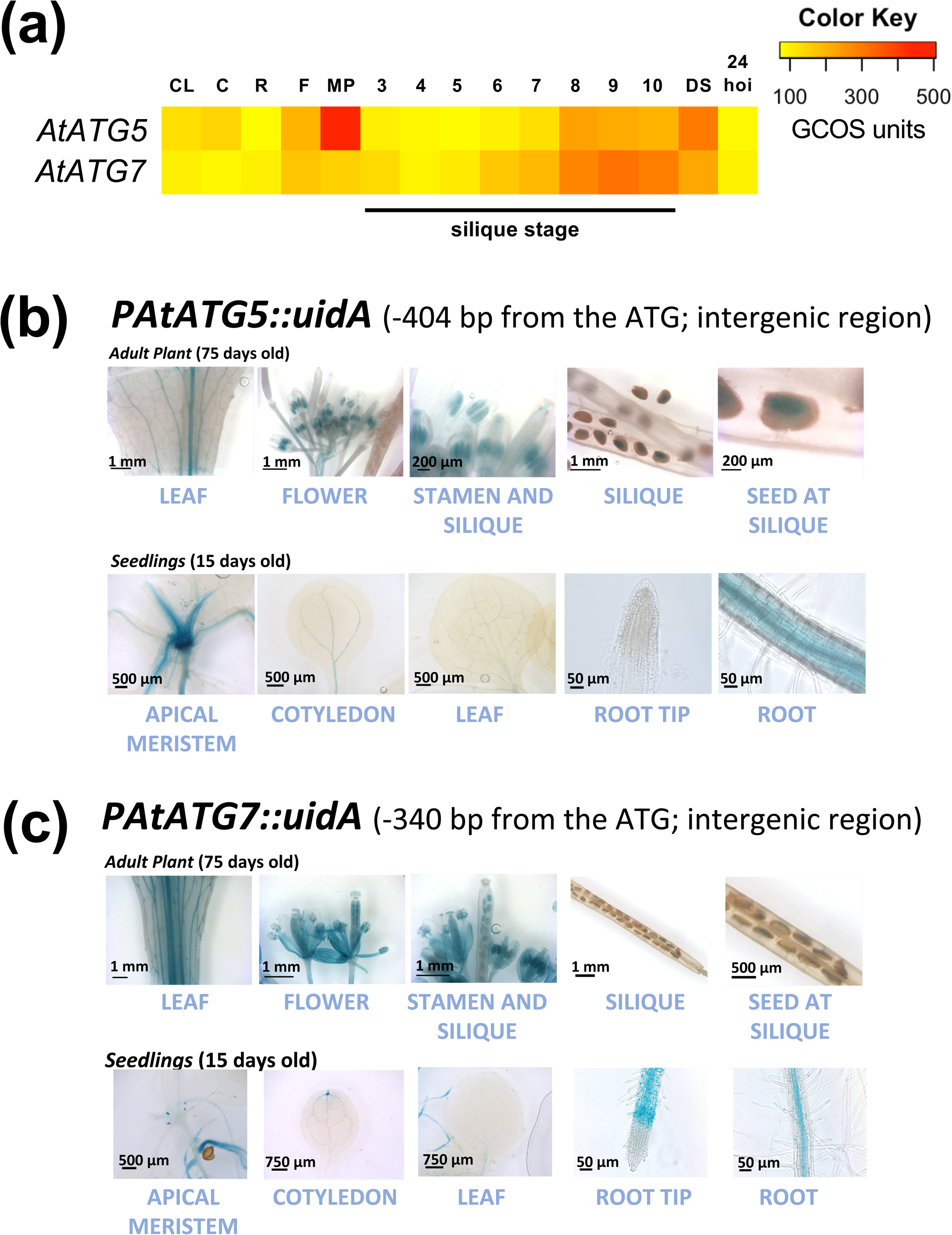
Temporal and spatial expression of *AtATG5* and *AtATG7* genes in adult plants. (a) Expression data of *AtATG5* and *AtATG7* were obtained from the Gene Expression Tool at the Bio-Analytic Resource for Plant Biology (BAR) public database and represented as a heatmap (scale 0-500 GCOS units). GUS histochemical staining of different organs and tissues from *PAtATG5::uidA* (b) and *PAtATG7: uidA* (c) transgenic adult plants. C: Cotyledon, CL: Cauline Leaf, DS: Dry Seed, F: Flower, R: Root, MP: Mature Pollen.

We further investigated the spatial expression patterns of *AtATG5* and *AtATG7* using stable transgenic Arabidopsis (Col-0) lines with *AtATG5* and *AtATG7* promoters driving GUS expression. Histochemical staining confirmed their ubiquitous expression in both adult plants (75 days) and seedlings (15 days), with GUS activity observed in the vascular elements of roots and leaves (Fig. 1b and 1c). Additionally, expression was also found in pollen grains and in seeds at late silique maturation stages. In seedlings, both genes were expressed in the stem apical meristem (SAM) and weakly in cotyledons (Fig. 1b and 1c).

Quantitative PCR (qPCR) was used to analyze *AtATG5* and *AtATG7* expression during embryogenesis, seed maturation, and germination, with *AtACT7* as a reference gene which showed stable expression throughout the period of study (Graeber *et al*., 2011; Fig. S2). Expression levels of both genes peaked during early green (EG) to late green (LG) and brown (B) stages of silique development, with maximal transcript accumulation in the B stage (Fig. 2b). During seed germination, *AtATG5* and *AtATG7* mRNAs were detected at high levels in the DS stage (Fig. 2c), followed by a sharp decline in the early stages of imbibition (12 hoi), and a slight increase between 24-30 hoi, when seeds reached t50 (time to 50% germination). GUS histochemical staining in *PAtATG5::uidA* and *PAtATG7::uidA* transgenic lines showed GUS activity at 24 and 48 hoi in the embryo but not in the endosperm. At 24 hoi, GUS activity was localized in the radicle transition zone, and by 48 hoi, the signal expanded to the hypocotyl and cotyledons, with increasing intensity as they developed (Fig. 2c). These findings suggest that *AtATG5* and *AtATG7* play important roles during seed maturation and germination.

**Fig. 2.**
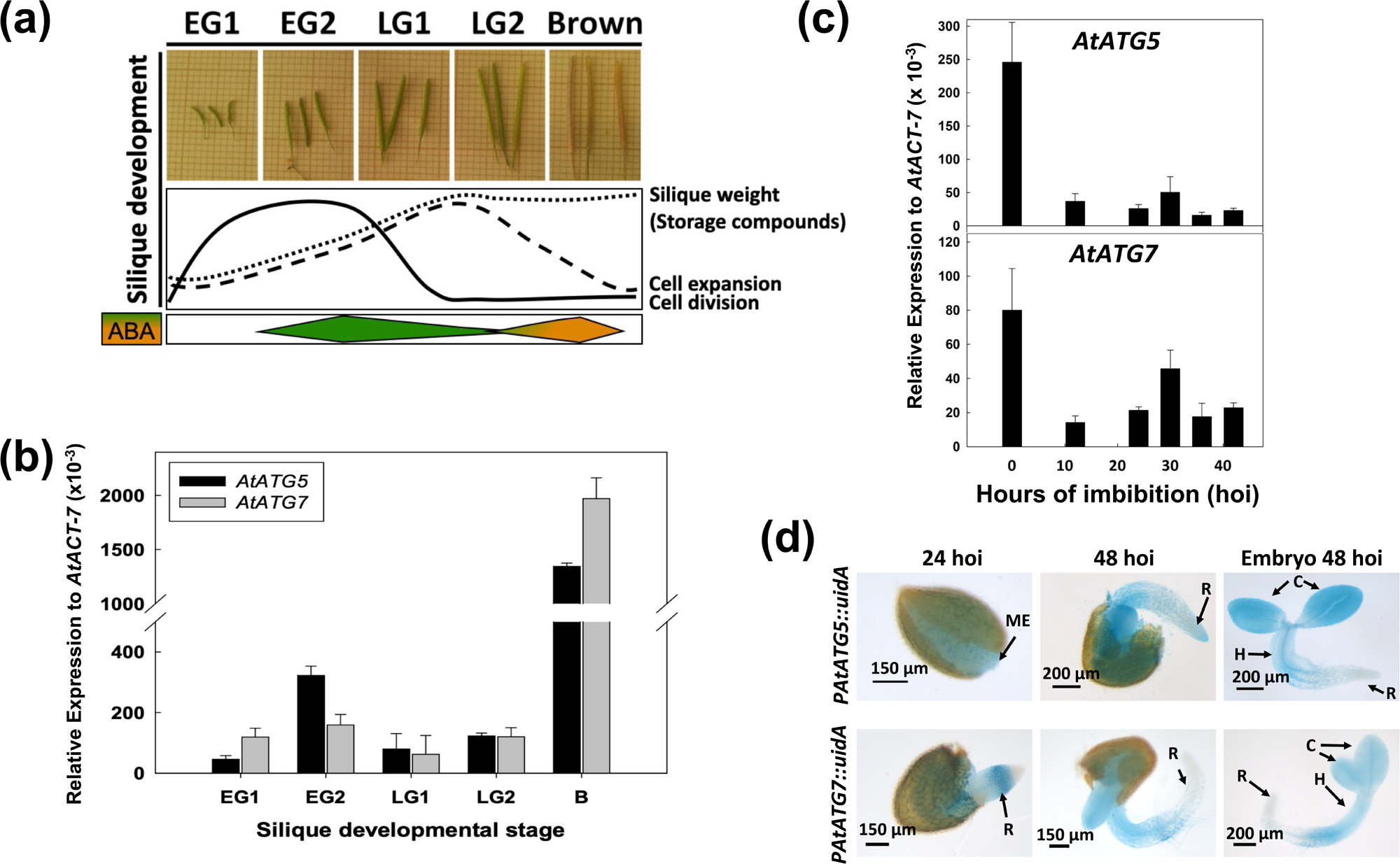
Expression patterns of *AtATG5* and *AtATG7* genes during embryogenesis, seed maturation and germination. (a) Images display the developmental stages of siliques used in the qPCR assays, along with corresponding patterns of silique weight, cell expansion, cell division, and ABA accumulation during silique development. Transcript analysis of *AtATG5* and *AtATG7* during seed embryogenesis and maturation is shown in (b), while (c) illustrates their expression during imbibition. (d) GUS histochemical staining of different *PAtATG5::uidA* and *PAtATG7::uidA* embryos at different time points (hours) of imbibition (hoi). Early Green stages 1 and 2 (EG1 and EG2), Late Green stages 1 and 2 (LG1 and LG2), and Brown (B) seeds. Data is shown as means ± standard error (SE) of three independent experiments, normalized to the expression of the *AtACT-7* gene. ABA peaks are shown in Fig. 2a, as described by Finkelstein *et al*. (2002, 2004).

### Seed germination is impaired and significantly delayed in *atg5* and *atg7* knockout mutants in the presence of ABA during imbibition

To investigate the relevance of *AtATG5* and *AtATG7* genes in seed germination, we analyzed homozygous T-DNA insertion mutant lines *atg5-1* and *atg7-2* (Hofius *et al*., 2009; Yoshimoto *et al*., 2009; Avin-Wittenberg *et al*., 2015; Barros *et al*., 2017; Erlichman *et al*., 2023) under both control conditions and in the presence of 1 μM ABA. The ABA-insensitive *abi5-7* mutant (Nambara *et al*., 2002; Albertos *et al*., 2015) was used as a negative control. The absence of *AtATG5* and *AtATG7* transcripts in the *atg5* and *atg7* mutants was confirmed by qPCR, while *abi5-7* exhibited basal expression (Fig. S3).

Seed Coat Rupture (SCR) and Endosperm Rupture were scored in Col-0, *atg5*, and *atg7* KO mutant lines (Fig. 3). Under control conditions, *atg5*, *atg7*, and *abi5-7* mutants showed a significant delay in germination, evidenced by an increased t50 for both SCR and ER compared to Col-0 (Fig. 3). While 1 μM ABA had minimal effect on Col-0 and *abi5-7* SCR, a notable delay in Endosperm Rupture was observed for *atg5* and *atg7* mutants, which did not reach t_50_ within the observed period. In contrast, Col-0 seeds reached t_50_ at 69.3 ± 4.2 h, while *abi5-7* seeds showed reduced ABA sensitivity, reaching t_50_ at 55.6 ± 1.2 h.

**Fig. 3.**
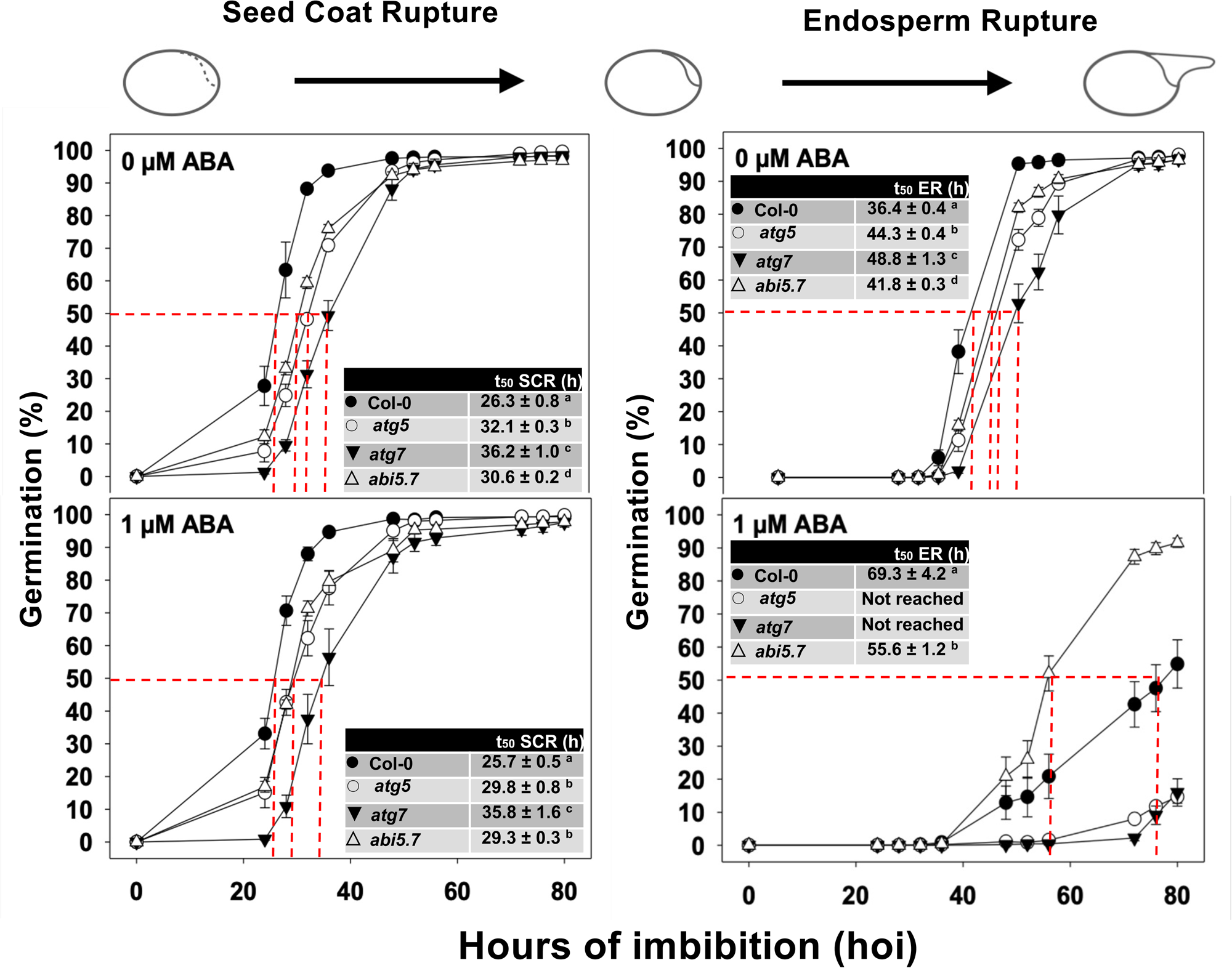
Germination assays in *atg5, atg7, abi5-7* mutant lines, and Col-0 (control) in the presence and absence of 1 µM ABA. Means ± SE of three independent replicates are represented. SCR: Seed Coat Rupture; ER: Endosperm Rupture, *sensu stricto* germination. In the inset, the time required to achieve 50% seed coat rupture (t_50_) and endosperm rupture (t_50_) is indicated.

Analysis of *PAtATG5::uidA* and *PAtATG7::uidA* imbibed seeds revealed that the spatial expression and intensity of GUS activity were not significantly altered in the presence of 1 μM ABA compared to control conditions (Fig. S4a). Additionally, RNA-seq analysis (FPKM; Fragments Per Kilobase Million, see next section) of *AtATG5* and *AtATG7* expression in the presence of ABA during imbibition showed results consistent with those observed under control conditions (Fig. S4b).

### Comprehensive transcriptomic profiling of *Arabidopsis thaliana atg7* mutant during seed imbibition reveals important changes in gene expression compared to the WT

To analyze transcriptional changes during seed germination in Col-0 and *atg7* mutants, under conditions with and without 1 μM ABA, RNA-seq analysis was performed on dry and imbibed seeds collected at 0, 24, 48, and 96 hours of imbibition (hoi) (Fig. S5a). These time-points correspond to t_0_, t_25_, t_75_, and t_100_, representing the time to reach 100% germination (Fig. S5a). Total RNA was extracted at each time point, and after library preparation, RNA sequencing was performed using the DESeq2 package in RStudio (see Materials and Methods). The raw DESeq2 output, including normalized counts, log_2_FoldChange, p-value, and adjusted p-value (padj; FDR corrected), is provided in Supplementary Data S2 and S4.

The hierarchical clustering analysis (Fig. S5b) revealed that the samples cluster primarily by their state (dry or imbibed) and secondarily by time points for imbibed samples. A clear distinction between mutant and WT samples was observed at each time point, especially for imbibed seeds. The principal component analysis (PCA) results (Figs. S5c-d) showed that the first Principal Component (PC1) explained most of the variance (78.2%), associated with differences between dry and imbibed samples. The second Principal Component (PC2) accounted for 15.9% of the variance and mainly reflected the time progression from 24h to 96h in imbibed seeds. Differences between the 48 hoi and 96 hoi time points may also be influenced by ABA treatment. Col-0 and *atg7* mutants were most clearly separated at 24 hours without ABA (Fig. S5c-d), and these conditions were chosen for differential gene expression analysis (DEG; Data S2).

The volcano plot of DEG (Fig. 4a) revealed higher expression of genes related to storage compound synthesis and mobilization in *atg7* mutant compared to WT. Gene set enrichment analysis (GSEA) (adjusted p-value < 0.05) identified ten pathways significantly induced and five significantly repressed in the mutant (Fig. 4b; Supplementary Data S3a). GO categories like lipid storage and seed maturation were overrepresented among induced categories, with lipid storage showing the highest Normalized Enrichment Score (NES). In contrast, response to ER stress (ERS), including autophagy, was repressed and showed the most negative NES. The repression of genes related to ERS and the induction of lipid storage genes suggests a clear trend in the mutant, with genes in these categories clustering at opposite ends of the list based on their Log_2_FC.

**Fig. 4.**
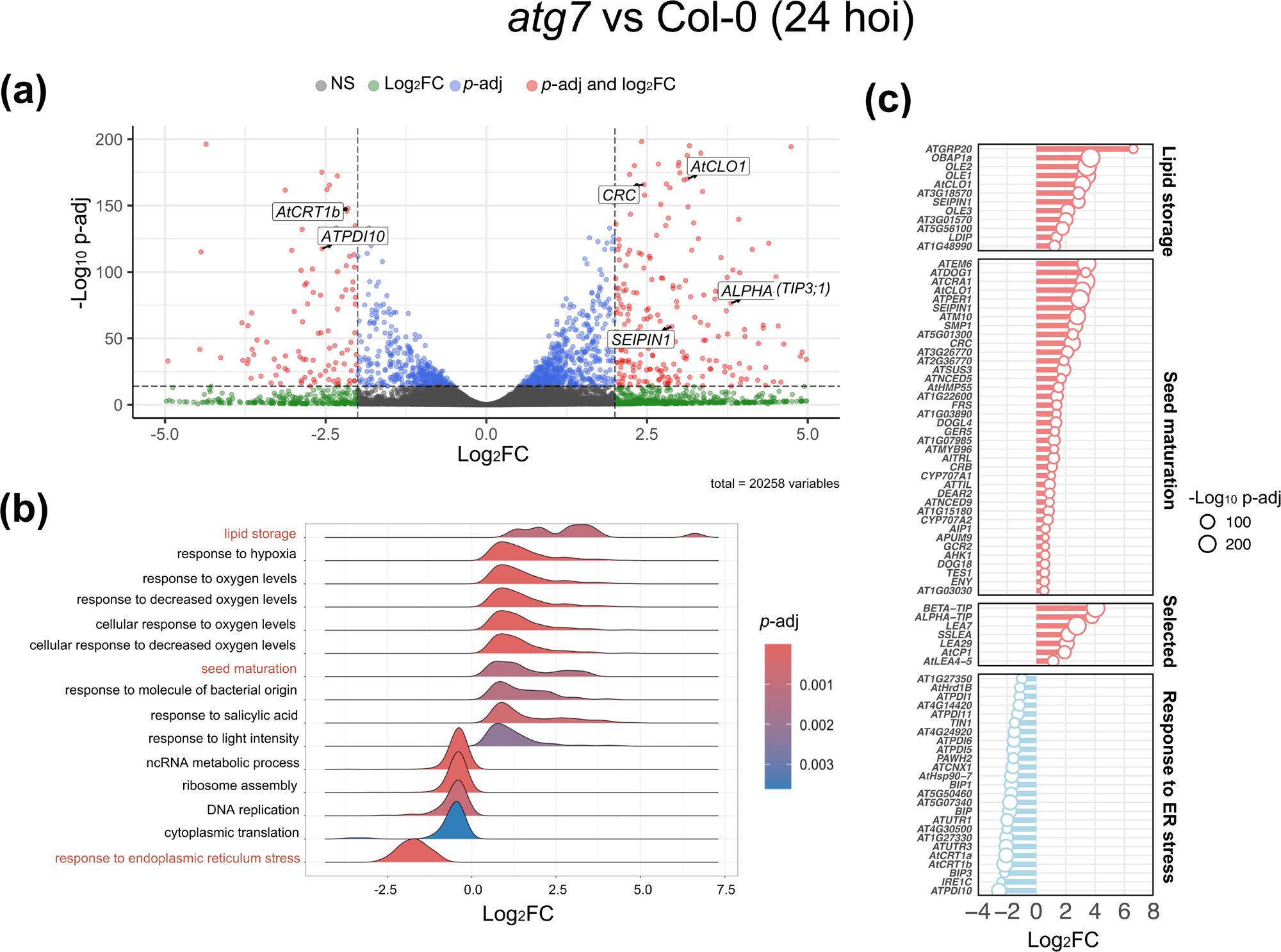
Differentially expressed genes (DEG) analysis in *atg7* in comparison to Col-0 at 24 hoi. (a) The volcano plot displays DEG as red dots, meeting the criteria of *p*-adj. ≤ 10e-16 and ± ≤ |2.| Some of the most significant DEGs such as *AtCLO1, CRC, ATPDI10, AtCRT1b, SEIPIN1, and ALPHA-TIP* are highlighted in boxes. (b) GSEA categorizes all DEG into different GOs represented as curves ranked according to their Log_2_FC. The categories highlighted in red are further analyzed. (c) The lollipop plot visualizes the expression of DEGs that meet the criteria in (a) and belong to the selected GO categories from (b). Additional DEGs that do not fall into the selected categories are included in the plot as well. NS, non-significant; Log_2_FC, Log_2_ fold change; *p*-adj, *p*-adjusted. *Indicate genes present in two categories.

Fig. 4c presents a lollipop plot illustrating the Log_2_FC of significant genes within selected categories (lipid storage, seed maturation, and ERS response). Genes involved in seed maturation, like *AtCRC* (*CRU3*), *AtCRA1* (*12S SSP*, cruciferins), *AtCLO1*, and *SEIPIN1*, were identified. *AtCLO1* encodes a caleosin protein involved in lipid body formation, while *SEIPIN1* is crucial for lipid droplet biogenesis in embryos. Additionally, lipid storage-related genes like *OLE1*, *OLE2*, and *OLE3* (oleosins) and *Late Embryogenesis Abundant* (*LEA*) protein-encoding genes (*SSLEA*, *LEA29*, *LEA7*) were also identified. Interestingly, two aquaporin-encoding genes, including *TIP3;1* (*ALPHA-TIP*), were differentially expressed. *TIP3;1* is notably the most expressed autophagy-related gene in seeds (Fig. S1).

As expected, genes related to ERS, including *BIP3* (HSP70 family) and *IRE1C* (involved in UPR in the ER), were repressed in the mutant (Fig. 4c). These findings suggest autophagy’s involvement in lipid and protein metabolism during early seed imbibition. Furthermore, 18 out of 38 genes in the seed maturation category were identified as potential ABI5 target genes through DAP-seq analysis (AGRIS Plant TF database linked to ShinyGO platform, Yilmaz *et al*., 2011; O’Malley *et al*., 2016; Ge *et al.,* 2020) (Data S3c).

Although other pathways like the response to oxygen levels and light were also identified as over-expressed in the mutant, lipid storage, ERS, and seed maturation categories were prioritized due to their significant differences and relevance in seed biology.

### Histochemical analysis of protein and lipids in *atg* mutants during seed imbibition reveals autophagy-related changes in the accumulation and dynamics of storage compounds

The transcriptomic analysis described above revealed a significant upregulation of genes associated with storage compounds in imbibed seeds of the *atg7* mutant compared to the WT (Fig. 4). To further investigate this finding, the detection of seed lipids and proteins was carried out in both *atg5* and *atg7* mutants, as well as in the WT, at 24 and 48 hoi.

Microscopic examination of embryos using Nile red for lipid staining (Fig. 5; Fig. S6: negative control) demonstrated similar levels in *atg5* and *atg7* mutants and Col-0 at 24 hoi (Fig. 5a). However, a quantitative analysis of the percentage of lipid /cell area in the radicle region for each line at 48 hoi (Fig. 5b) showed that the percentage of lipid/cell area is significantly lower in mutants (*ca.* 30%) than in WT (ca. 45%). The *atg7* mutant exhibits significantly reduced expression of 15 genes related to fatty acid biosynthesis, in addition to 2 genes specifically involved in fatty acid desaturation and 5 genes associated with fatty acid elongation, compared to Col-0 at 24 hours (Figure S7a, Data S3d). These genes encode enzymes that catalyze 7 reactions involved in fatty acid synthesis and 1 reaction in the extension of palmitoyl-CoA and stearoyl-CoA into very-long-chain acyl-CoAs (KEGG; Figure S7b, c). Interestingly, three genes encoding lipases related to lipid catabolism (*MPL1*, *MAGL4*, and *MAGL12*) as well as *ACX1*, which encodes the first enzyme in the β-oxidation pathway, are significantly induced in the *atg7* mutant compared to the WT during imbibition (Figure S7).

**Fig. 5.**
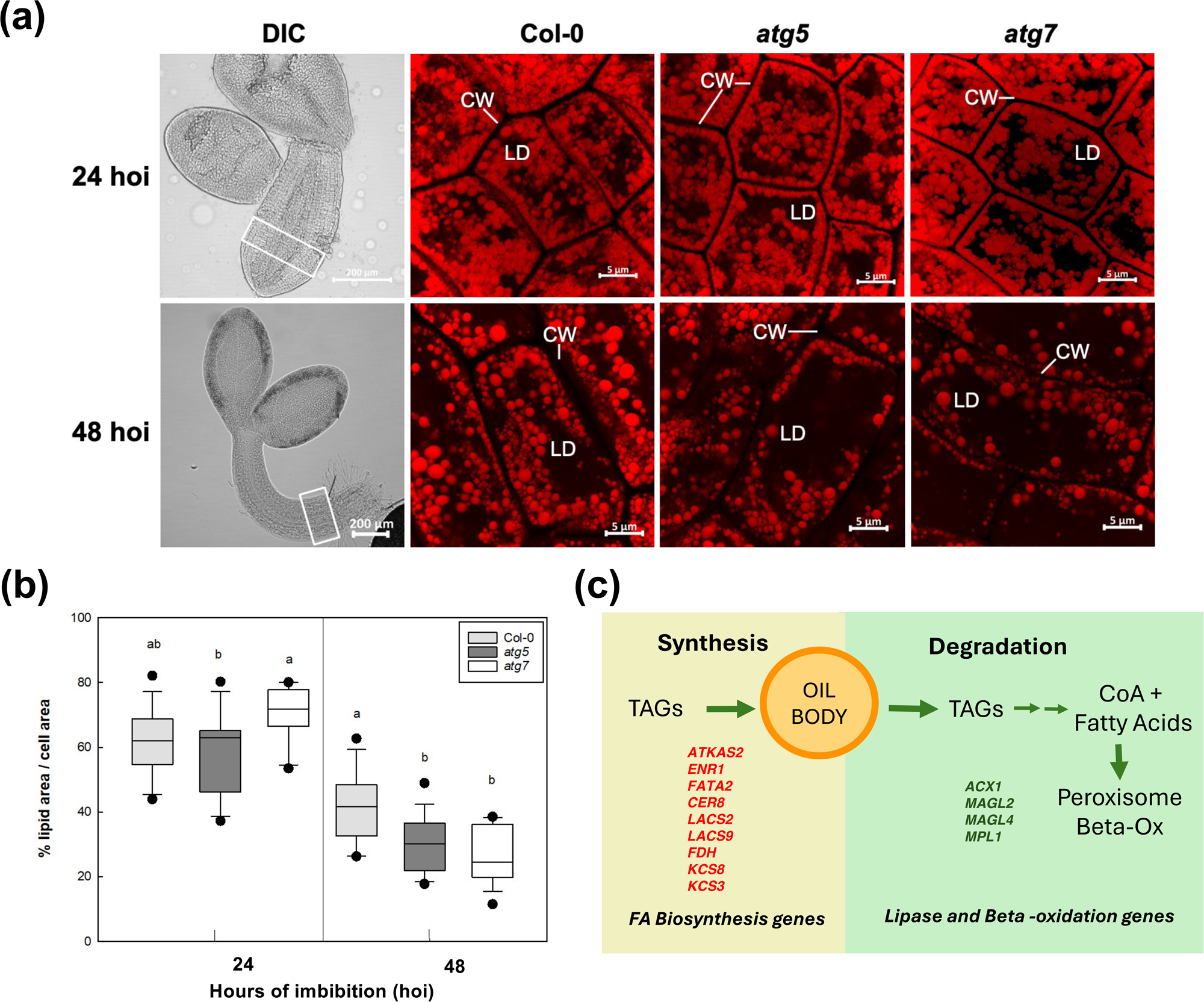
Differences in lipid content in *A. thaliana* Col-0, *atg5*, and *atg7* embryos at 24 and 48 hoi. (a) Differential Interference Contrast (DIC) images at x10 magnification show the entire embryos, with the radicle area used for lipid visualization and lipid content analysis marked with a rectangle. Confocal images of Nile Red lipid staining of individual embryo cells are visualized at x60 magnification. (b) Lipid content analysis of individual cells represented as % lipid area/cell area in the different *A. thaliana* lines. Data shown are means ± SE of at least 10 independent cells. Letters indicate statistically significant differences among the three Arabidopsis lines (p≤0.05, One-way ANOVA followed by Turkey’s comparisons test). (c) Dynamics of lipid accumulation in seed embryos. The depicted scheme shows the balance between oil body biogenesis and degradation, indicating participating enzymes with altered gene expression levels in the *atg7* mutant at 24 hoi (red: repressed; green induced), see Fig. S7 for more details.

Likewise, our analyses revealed differences when *atg7* was compared with WT in the ontology term of genes related to seed maturation proteins. Examples include *AtCRC* (*CRU3*) and *AtCRA1*, which encode *12S-SSP* (cruciferins), highly abundant SSP in seeds of *A. thaliana.* These proteins accumulate in PSV during seed maturation. To investigate SSP in Col-0, *atg5*, and *atg7* seeds, in DS and imbibed stages, a histochemical analysis was performed. Longitudinal sections of *A. thaliana* seeds (DS and water-imbibed at 24 and 48 hoi) were stained using Periodic acid–Schiff (PAS) and Naphthol Blue Black to detect insoluble polysaccharides (mainly cellulose and starch) and proteins, respectively. As shown in Fig. 6a–f, both DS and imbibed seeds exhibited abundant proteins (blue stained). In Col-0 DS, proteins accumulated within well-defined PSV are characterized by rounded structures (Fig. 6a-b). On the contrary, both *atg5* and *atg7* mutants, although rich in proteins as well, exhibited altered organization in their cytoplasm, as the rounded structures seen in WT embryo cells were not clearly distinguished, especially in the radicle (Fig. 6a-r). These differences in cytoplasmic organization, relative to protein accumulation in radicle cells of *atg* mutants, persist at 24 hoi (Fig. 6g-l). At 48 hoi, proteins were completely degraded in both Col-0 and mutants after radicle emergence (Fig. 6m–r).

**Fig. 6.**
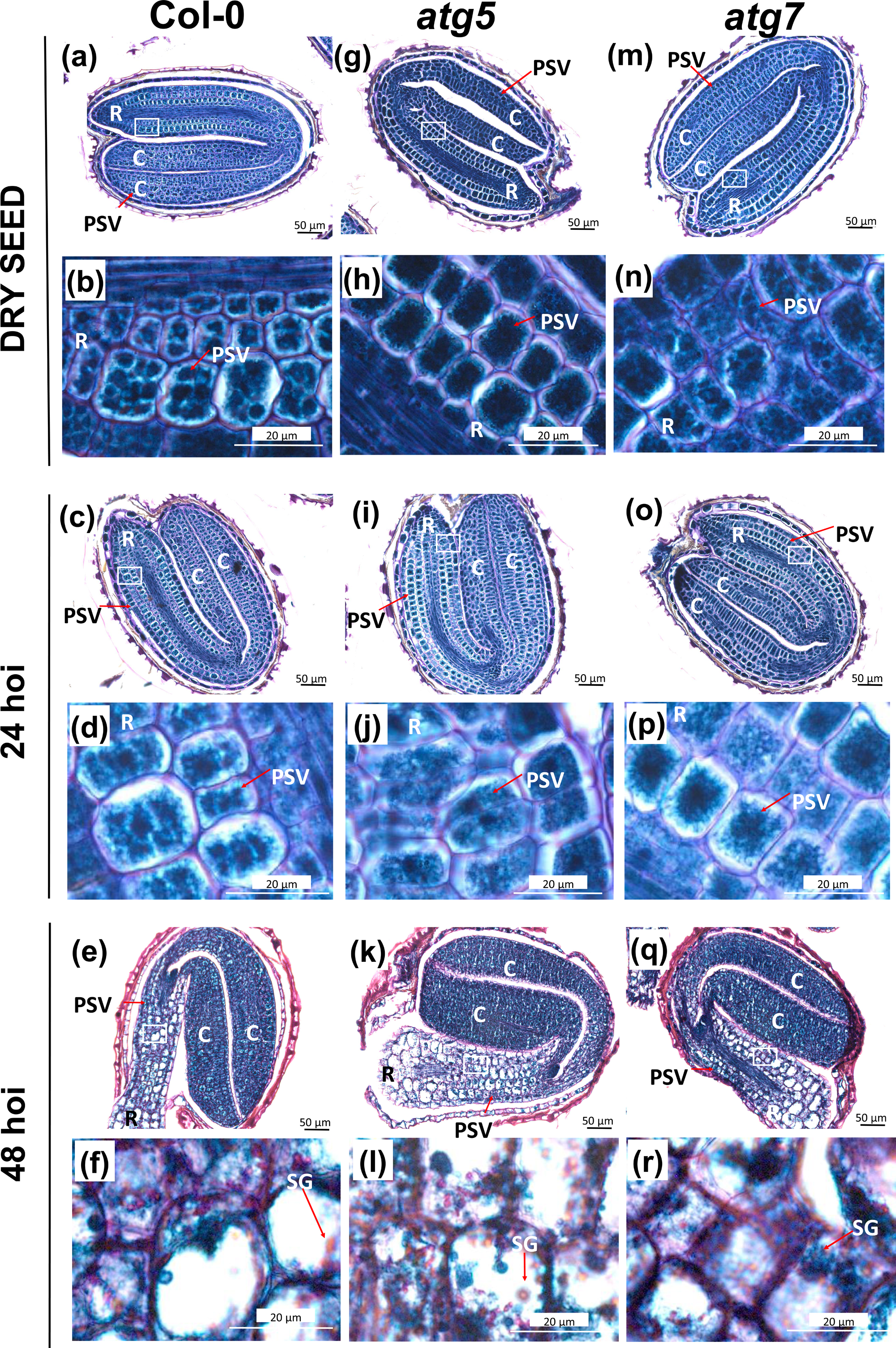
Differences in protein organization in *A. thaliana* Col-0, *atg5*, and *atg7* dry (0 hoi) and 24 and 48 hoi seed sections. Periodic acid (PAS)-Naphthol Blue Black (NBB) staining of protein and polysaccharides is shown for Col-0 (a-f), *atg5* (g-l), and *atg7* (m-r) seed transversal sections. Images of DS (a, g, m), at 24 hoi (c, i, o) and at 48 hoi (e, k, q) taken at 25x magnification, showing with a white rectangle indicating the close-up image taken at 60x magnification for DS (b, h, n), at 24 hoi (d, j, p) and 48 hoi (f, l, r). C, cotyledon; PSV, Protein Storage Vacuoles; R, radicle; SG, starch grain.

### Immunolocalization of the ATG8 protein in PSVs of WT but not in *atg* mutants supports an autophagy-like mechanism involved in reserve mobilization during seed germination

The insights presented above from genetic, transcriptomic, and histochemical analyses during seed imbibition and reserve mobilization prompted us to explore the molecular link between storage compound mobilization and autophagy. To this end, we performed immunofluorescence labeling using a primary antibody specific to ATG8a protein (Miklaszewska *et al*., 2023) participating in autophagosome capture of cargoes (Fig. 7a-l, Fig. S8). The presence of the ATG8a protein was detected in longitudinal sections of Col-0 seeds imbibed at 24 hoi. As illustrated in Figs. 7a-c, ATG8a is predominantly localized within PSVs in the radicle of Col-0 seeds. Interestingly, this localization was practically absent in *atg* mutants, specially in*atg7* embryos (Fig. 7e-g, i-k). Secondary/primary antibodies and empty negative controls did not show fluorescence signals as expected (Figs. S8a-i). Examining coarse sections (8 μm) of paraffin samples through differential interference contrast microscopic analysis revealed that the cellular organization of seed tissues had been well preserved (Fig. 7d, h, l; Fig. S8c, f, i). Although these data do not rule out the presence of other ATG8 proteins in PSVs in the radicle of *atg5* and *atg7* embryos, our histochemical assays support the participation of autophagy not only in lipid and protein metabolism, as suggested by our transcriptomic data, but also in its cytoplasmic organization of embryo cells throughout seed germination.

**Fig. 7.**
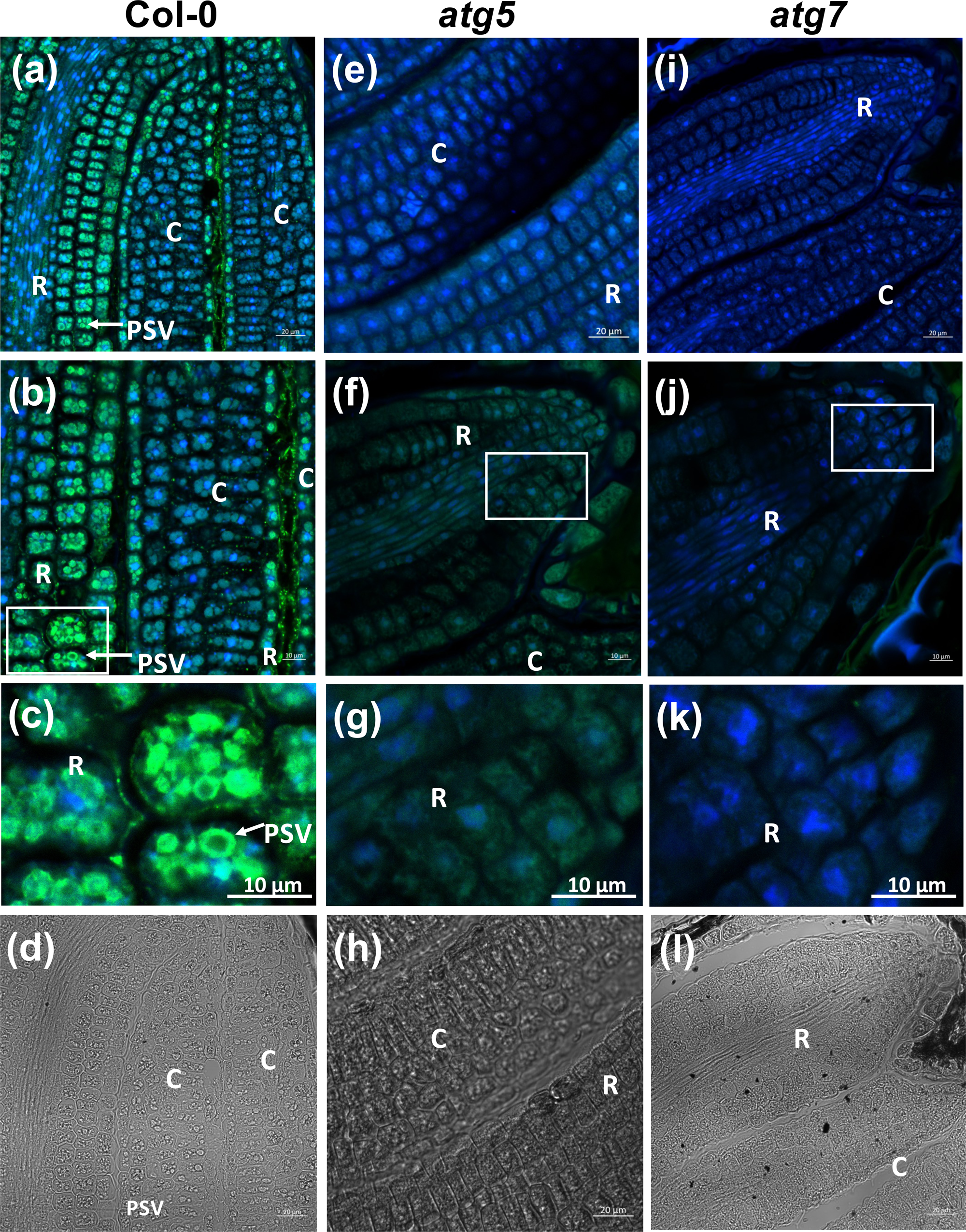
Differences in PSV organization in *A. thaliana* Col-0, *atg5*, and *atg7* germinating seed sections. Confocal images showing AtATG8 immunolocalization in Col-0 (a-c), *atg5* (e-g), and *atg7* (i-k) seed transversal sections at 40x magnification (a, e, i), 63x magnification (b, f, j) and a close-up image from x63 magnification, indicated with a white rectangle (c, g, k). DIC images at x40 magnification are also shown (d, h, l). C, cotyledon; PSV, Protein Storage Vacuoles; R, radicle.

### Transcriptomic profiling of *atg7* imbibed seeds reveals a connection of autophagy to ABA responses and genes regulated by ABI5

ABA is a hormone that regulates seed development and germination. The autophagy mutants exhibited delayed germination compared to WT, particularly with 1 μM ABA during imbibition (Fig. 3). To explore the molecular basis of ABA responses in the *atg7* mutant, we compared transcriptional differences during seed imbibition in the absence and presence of ABA between Col-0 and the *atg7* mutant (Fig. S5). The ABA treatment at 24 hoi had no significant effect (Fig. S5). Therefore, we compared the transcriptomes of germinating *atg7* and Col-0 seeds at 96 hoi with ABA and 48 hoi without ABA (Figs. 4, S5; Data S4 and S5). For differential expression analysis, we used the model: ∼ Treatment + Mutant + Treatment:Mutant, and performed a likelihood ratio test against the reduced model ∼ Treatment + Mutant to assess the interaction term’s significance. Genes were considered significantly differentially expressed when the adjusted p-value was < 0.05 and the log_2_fold change (log_2_FC) was ≥ ±1.

In the volcano plot (Fig. 8a; Supplementary Data S4), DEGs show whether the effect of ABA differs between *atg7* and Col-0. A positive Log_2_FC represents genes where ABA effect is more pronounced in the mutant (“enhanced ABA effect genes”; n = 308), and a negative Log_2_FC represents genes with a reduced ABA effect in the mutant (“reduced ABA effect genes”; n = 885). Several ABA-responsive genes, such as *EM1* and *AFP1* (Bensmihen *et al*., 2002; López-Molina *et al*., 2003; Du *et al*., 2024), are highlighted.

**Fig. 8.**
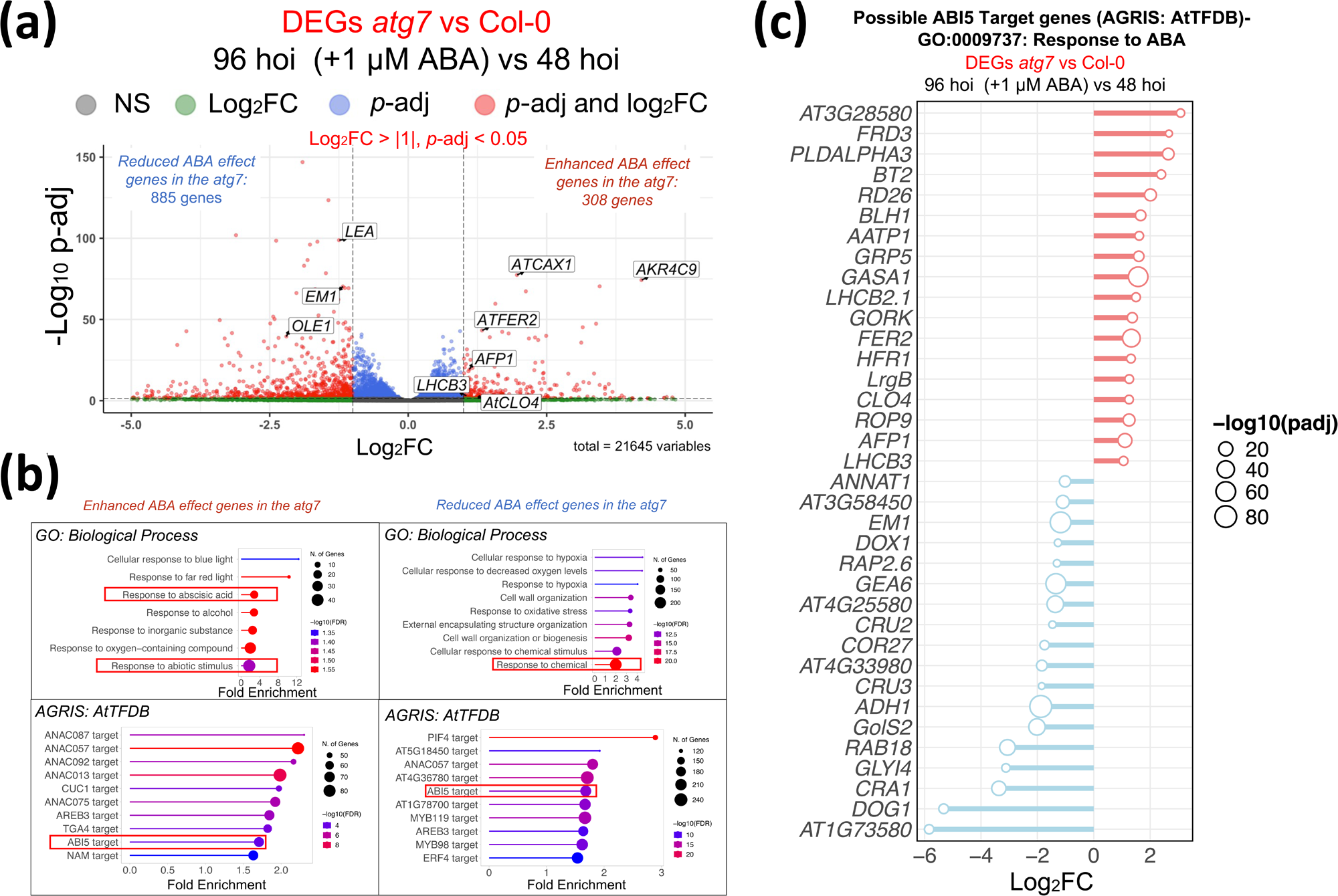
Transcriptomic analysis of the specific ABA response in *atg7* and Col-0 imbibed seeds. (a) The volcano plot displays DEG as red dots, meeting the criteria of *p*-adj < 0.05 and Log_2_FC ≥ |2.0|. Some of the most significant DEGs such as *AFP1, AtCAX1, AtCLO4, AtFER2, EM1, and OLE1,* are highlighted in boxes. (b) Gene ontology (GO) analysis using ShinyGO platform of DEG in (a), key gene categories are marked in red, focusing on ABA and abiotic stress-related categories (top plots). Transcription factor (TF) analysis using the AGRIS Plant TF database linked to ShinyGO platform for potential regulators of the DEG in (a; bottom plots). (c) The lollipop plot visualizes the expression of DEGs identified as possible ABI5 targets and belong to ABA response GO term in (b). hoi, hours of imbibition; NS, non-significant; Log_2_FC, Log_2_ fold change; *p*-adj, *p*-adjusted. ABI5 target genes extracted from AGRIS Plant TF database are reported in Besmihen *et al*. (2002) and O’Malley *et al*. (2016).

GO term analysis (ShinyGO platform) revealed significant enrichment of genes related to the “response to chemical” GO term in the *atg7*-reduced gene group (Fig. 8b, top right panel). In contrast, the *atg7*-enhanced gene group showed enrichment for “response to abiotic stimulus,” particularly ABA response (Fig. 8b, top left panel).

268 of the 1,193 (22.5%) DEGs identified were previously identified as potential *ABI5* target genes (Fig. 8b, bottom panels; Supplementary Data S5c and S5d). Of these, 36 genes were classified in the ABA response category (Fig. 8c; Supplementary Data S5e, S5f). Fig. 8C presents a lollipop plot illustrating the Log_2_FC of these 36 genes (adjusted p-value < 0.05).

### The autophagy protein ATG8 directly interacts with the TF ABI5 both *in vivo* and *in vitro*, supporting its modulation through autophagy

The transcriptomic analysis previously described supported the role of ABA in seed development and suggested the connection of the ABI5/ABI3 signalling to autophagy during seed germination. To deepen our findings linking these TFs with the autophagy machinery, we focused on ABI5, a bZIP TF primarily expressed in seeds and associated with suppression of germination and post-germination growth under adverse conditions (López-Molina *et al*., 2001; Albertos *et al*., 2015). ABI5 accumulation was systematically monitored in the *atg5* and *atg7* mutants, both in the DS and during imbibition, compared to the WT seeds. Western blot assays were performed at 0, 36, 48, and 72 hoi using a primary antibody specific for the ABI5 protein (Fig. 9a). The signal intensities of the ABI5 multiple forms in the Western blot assays were quantified using ImageJ software and plotted in Fig. 9b. This Fig. shows that multiple forms of the ABI5 are clearly detected in WT seeds in DS (0 hoi) but shows increased levels in autophagy mutants *atg5* (nearly 2-fold) and *atg7* (about 2.5-fold). This difference in ABI5 accumulation was further monitored at different time points during imbibition (Figs. 9a and 9b) where it shows a temporal decline until untraceable levels around 72 hoi. These findings indicate that autophagy can be important in modulating ABI5 accumulation during seed imbibition. Furthermore, we verified that another bZIP factor expressed during seed development, bZIP67. This TF is the closest homolog of ABI5, and was previously described as a regulator of omega-3 fatty acid content in Arabidopsis seeds (Mendes *et al*., 2013; Sánchez-Vicente *et al*., 2024).bZIP67 also accumulates to a greater extent in autophagy mutants *atg5* and *atg7* and shows decreased levels in *ATG5* and *ATG7* overexpression lines (Fig. S9a). Moreover, bZIP67 contains three possible ATG8-interacting motifs (AIMs) within its sequence (Fig. S9b). Interestingly, genes such as *cruciferin 1* (*CRU1*), *CRU2*, *CRU3*, *seed storage albumin 1* (*AT2S1*) and *fatty acid elongation 1* (*FAE1*), previously recognized as bZIP67 target genes with roles in protein and lipid metabolism in seeds (Mendes *et al*., 2013), were enhanced by ABA in *atg7*-imbibed seeds compared to Col-0, as revealed by our transcriptomic analysis (Fig. S9c).

**Fig. 9.**
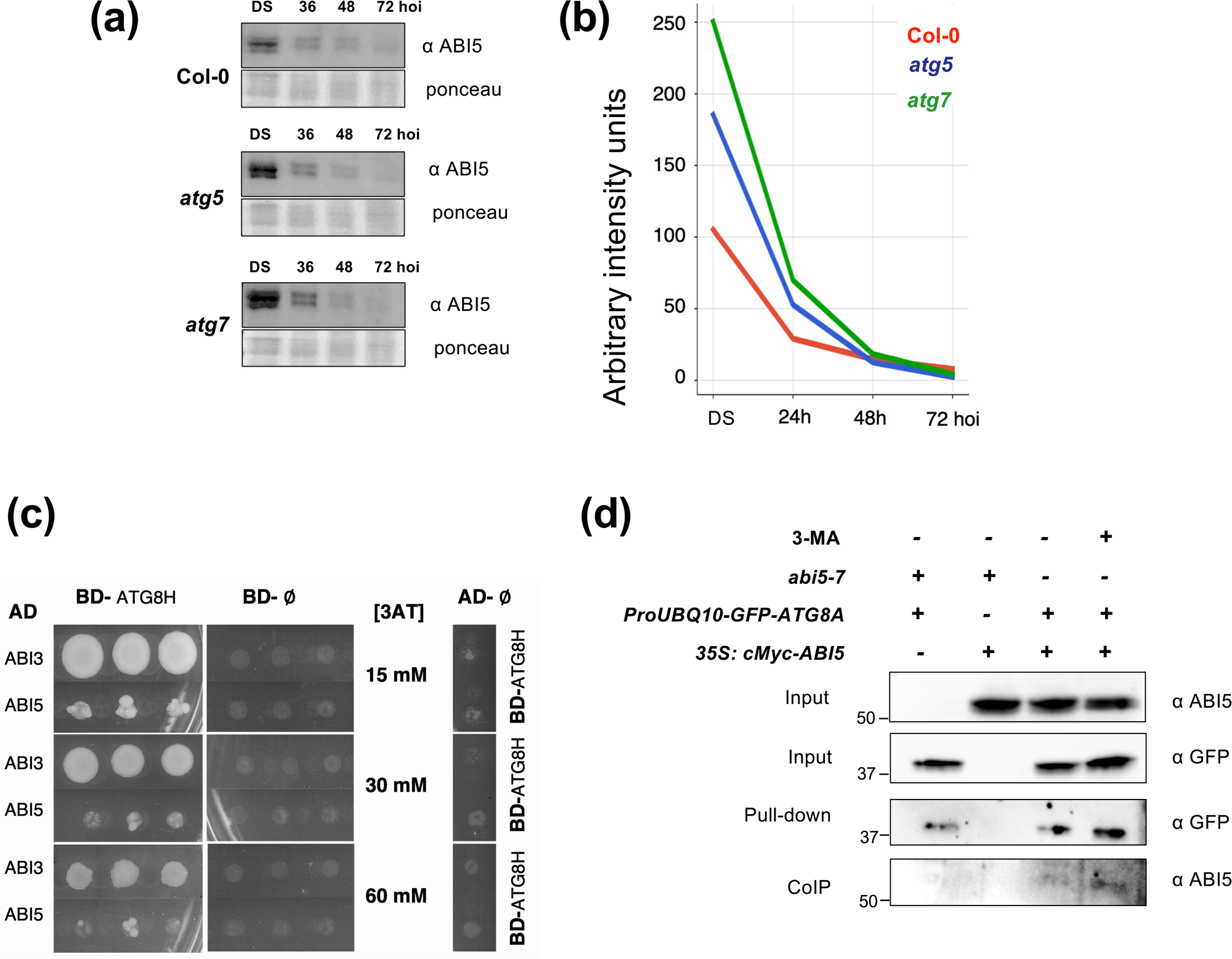
AtATG8H specifically binds to AtABI3 and AtABI5 proteins in a Y2H experiment and AtABI5 degradation is delayed in *atg5* and *atg7* mutants. (a) Western blot analysis of AtABI5 in *A. thaliana* Col-0, *atg5* and *atg7* dry (0 hoi) and germinating seeds (36, 48, 72 hoi). (b) The ABI5 band intensity from western blot analysis in (b) is calculated by densitometry (means of the band found in each sample). (c) Growth of yeast diploid cells expressing the GAL4 binding domain fused to ATG8H (BD-ATG8H) and the GAL4 activation domain fused to ABI3 or ABI5 (AD-ABI3 or AD-ABI5) in selective medium (–L, –W, –H) with increasing concentrations of 3-AT. The lack of growth of the negative controls BD-lll and AD-lll is also displayed. (d) Interaction of ATG8-GFP and cMyc-ABI5. CoIP assays between *35S:cMyc-ABI5* and *ProUBQ10-GFP-ATG8A* Arabidopsis 4-day-old seedling extracts. Plants were grown in MS and the treated or not with 3-MA for 24 hours. Inputs levels were determined with anti-GFP and anti-ABI5 antibodies. Pull-down was revealed with anti-GFP antibody and the CoIP was revealed with anti-ABI5.

ABI5 serves as a master regulator in ABA signaling cascades within seeds as part of a complex with ABI3 (a B3 TF). These proteins play critical roles as seed-specific downstream ABA targets and are recognized as key repressors of germination (Sajeev *et al*., 2024). Sequence analysis revealed that both ABI3 and ABI5 contain several AIMs, with 6 and 5 potential AIMs, respectively (Fig. S10). This raised the question of whether ABI3 and ABI5 could physically interact with components of the autophagy machinery, particularly ATG8, during seed germination. To investigate this possibility, we examined their interaction with ATG8 using yeast two-hybrid (Y2H) assays (Fig. 9c). We focused on the ATG8H protein, the major ATG8 gene expressed during seed germination (https://bar.utoronto.ca/) and fused its coding sequence to the GAL4 DNA binding domain as bait. Yeast cells expressing the BD-ATG8H fusion along with either AD-ABI3 or AD-ABI5 were able to grow in selective medium containing more than 60 mM 3-aminotriazole (3-AT). This result, in contrast to the lack of growth in control cells or those expressing only AD-ATG8H, indicates that both ABI3 and ABI5 can indeed interact with ATG8 *in vitro* (Fig. 9c). With this information, we checked again the interaction by *in vivo* CoIP assays using the *35S:cMyc-ABI5* and *ProUBQ10-GFP-ATG8A* lines (Fig. 9d). Protein extracts from 4-day old seedlings, treated or not with the autophagy inhibitor (3-MA, 5 mM),were mixed and analyzed. The *GFP-ATG8A* protein was selected as bait and immunoprecipitated with anti-GFP beads, and the presence of *cMyc-ABI5* was detected by the anti-ABI5 antibody. Interestingly, the ABI5-ATG8A interaction was detected mainly in the presence of 3-MA, indicating that autophagy may control the ABI5 abundance during early stages of development.

## Discussion

In recent years, autophagy has been increasingly linked to a diverse array of plant biological processes, including development, nutrient recycling, and responses to both abiotic and biotic stresses. Although several studies have acknowledged its role in seed development (e.g., Michaeli *et al*. 2016; Minina *et al*., 2018; Tarnowski *et al*., 2020), the molecular mechanisms and targets involved remain partially understood. In this work, our aim was to explore its impact on seed germination in *A. thaliana* through the study of autophagy mutants *atg5* and *atg7* with a special focus on their connections to ABA signaling.

### Autophagy mutants *atg5* and *atg7* show a delayed seed germination phenotype and hypersensitivity to ABA

*AtATG5* and *AtATG7* are single-copy genes in the Arabidopsis genome, and their gain- and loss-of-function mutants have been used to study autophagy-related phenotypes (Hofius *et al*., 2009; Yoshimoto *et al*., 2009; Minina *et al*., 2018; Ohlsson *et al*., 2023). To investigate their role in germination, we examined their expression patterns during seed maturation and germination. Transcriptomic data from the BAR database showed increased expression of both genes in dry seeds, early imbibed seeds, late silique maturation, and mature pollen. These results were confirmed by qPCR and GUS histochemical staining (Fig. 1 and 2). The *ATG5* and *ATG7* promoters drove expression in vascular elements of roots, leaves, mature pollen, and seeds in late silique maturation, consistent with previous reports in Arabidopsis and other species (Thompson *et al*., 2005; Li *et al*., 2015; Yu and Hua, 2022). Expression was also observed in the stem apical meristem and cotyledons. Autophagy, associated with stress-enhanced microspore embryogenesis, plays a role in pollen germination, tube growth, and sperm cell development (Yan *et al*., 2023; Testillano, 2019; Berenguer *et al*., 2021), and is involved in forming xylem and phloem elements (Michalak *et al*., 2024).

Our results support earlier studies showing increased expression of *ATG* genes during seed development, especially during maturation (Di Berardino *et al*., 2018; Bedu *et al*., 2020). *atg5* and *atg7* mutants exhibit reduced nitrogen content, possibly due to defective nitrogen remobilization. Evolutionary conservation of ATG function was also observed in species like *M. truncatula* and *Zea mays* (Guiboileau *et al*., 2012; Li *et al*., 2015; Barros *et al*., 2017; Minina *et al*., 2018; Sera *et al*., 2019; Liu *et al*., 2023). *AtATG5* and *AtATG7* expression peaks during late seed maturation (Fig. 2a), coinciding with ABA increase, storage compound synthesis, and primary dormancy induction (Ali *et al*., 2021). These findings led us to investigate the germination capacity of *atg5* and *atg7* seeds. Germination assays showed significant delays in mutants compared to WT, particularly under ABA treatment. Slower germination in *atg* mutants was reflected by increased t_50_ values (Fig. 3).

In transgenic seeds expressing *uidA* under the control of the *AtATG5* and *AtATG7* promoters, GUS activity was observed in the radicle transition zone at 24 hoi, especially for *AtATG7*, and extended to the hypocotyl and cotyledons at 48 hoi. This pattern aligns with reserve mobilization involving SSP hydrolysis first in the embryonic axis and then in cotyledons (Müntz, 2007; Iglesias-Fernández *et al*., 2014). These results suggest that *AtATG5* and *AtATG7* influence germination by controlling storage mobilization in the radicle. Interestingly, F1 seeds from maternal *atg* mutants germinated earlier, likely due to increased water permeability and normal autophagy in the embryo. Maternal *atg* mutant seeds showed reduced mucilage content and structural changes in seed coat analysis (Erlichman *et al*., 2023).

In the presence of ABA, *atg5* and *atg7* mutants exhibited slower germination compared to WT. However, the spatiotemporal expression of *AtATG5* and *AtATG7* was unchanged by ABA in the seed imbibition medium (Fig. S4), suggesting ABA does not affect their expression but alters other factors impacting germination. Our data suggest a role for autophagy in seed germination progression under both non-stress and ABA-induced stress conditions.

### Transcriptomic analysis and reserve mobilization in *atg7* mutant seeds during imbibition

We compared the transcriptomes of WT and *atg7* seeds at 24 hoi to identify autophagy-related changes. DEGs analysis revealed significant transcriptional differences, with GO enrichment in processes linked to storage compounds, especially lipids and proteins. We thus performed histochemical analyses of protein bodies and lipid droplets in dry and imbibed seeds.

In *A. thaliana*, lipids are stored as TAG in lipid droplets (LD) stabilized by proteins like oleosins and caleosins (Miklaszewska *et al*., 2023). *AtOLE1* and *AtOLE2* are key seed oleosins for LD formation and degradation (Simada *et al*., 2008; Chen *et al*., 2019; Shao *et al*., 2019). At 24 hoi, *atg7* seeds show increased expression of *AtOLE1*, *AtOLE2*, and *AtOLE3*, suggesting a compensatory mechanism. Overexpression or suppression of oleosins delays germination, confirming their role (*Siloto et al.*, 2006; Yuan *et al*., 2021). DEGs also include *AtSEIPIN*, involved in LD biogenesis (Zhao *et al*., 2023), and *AtCLO1*, required for LD degradation and LD-vacuole interaction (Naested *et al*., 2000; Kim *et al*., 2011; Poxleitner *et al*., 2006). Interactions between *AtATG8* and both *AtCLO1* and *AtOLE1* via AIMs support autophagy’s role in LD dynamics (Marshall *et al*., 2019; Miklaszewska *et al*., 2023). The upregulation of LD-structural protein genes in *atg7* seeds supports autophagy’s involvement in lipid mobilization.

Autophagy mutants (*atg5, atg7*) exhibit reduced TAG and FA levels (Minina *et al*., 2018). Fan et al. (2019) showed that this reduction stems from impaired lipid recycling, not synthesis. Havé et al. (2019) reported altered phospholipid/sphingolipid accumulation in *atg5*, affecting ER and plasma membranes. Consistent with these, our Fig. 5 shows that while LD numbers in *atg7* seeds at 24 hoi are similar to WT, lipid degradation is faster at 48 hoi, highlighting a role for autophagy in early lipid mobilization.

The *atg7* mutant also shows reduced expression of FA biosynthesis, desaturation, and elongation genes, confirming autophagy’s role in lipid homeostasis (Bouchnak *et al*., 2023). Conversely, genes related to FA catabolism, such as *MPL1*, *MAGL4*, *MAGL12*, and *ACX1* (first β-oxidation enzyme), are upregulated. Genes like *CER8*, *LACS2*, and *LACS9*, traditionally linked to lipid synthesis, also participate in catabolism (Bai *et al*., 2021). Our RNA-seq at 24 hoi reflects early transcriptional changes corresponding to lipid shifts observed at 48 hoi (Fig. 5a), in line with Barros *et al*. (2021). Together, these suggest a shift toward lipid breakdown in *atg7*.

Arabidopsis seeds store 12S globulins (cruciferins) and 2S albumins (napins) in PSVs (Baud *et al*., 2008). *AtCRU1* and *AtCRU3* transcripts are elevated in *atg7* at 24 hoi. Histochemical analysis at DS and 24 hoi shows PSV disorganization in the radicle of *atg5* and *atg7* seeds, which resolves by 48 hoi. This suggests two protein trafficking routes to PSVs: Golgi-dependent and autophagy-like ERvt (Hills, 2004; Vitale *et al*., 2022). ATG8 immunolocalization in Col-0 at 24 hoi confirms its presence in PSVs, while it is absent in *atg* mutants, indicating impaired ERvt trafficking and protein degradation in the embryonic axis.

### A connection of the delayed germination phenotype and ABA responses to the stability of ABI5 in the *atg7* mutant

ABA is a key phytohormone regulating seed maturation, dormancy, and stress responses (Ali *et al*., 2021). We found that *atg5* and *atg7* mutants exhibit delayed germination (Fig. 3), especially under ABA treatment. Transcriptomic analysis comparing *atg7* and Col-0 with/without ABA revealed significant DEGs, with ABA response genes notably enriched among upregulated genes in *atg7*. About 22.4% of these DEGs are predicted ABI5 targets, suggesting altered ABA signaling.

ABI5, together with ABI3, regulates ABA-responsive genes suppressing germination and controlling preharvest sprouting (Koornneef *et al*., 1984; Finkelstein, 1994; Abraham *et al*., 2016; Matilla, 2022; Sajeev *et al*., 2024). ABA modulates germination based on environmental cues (López-Molina *et al*., 2001). A connection between ABA signaling and autophagy involves NBR1, a selective autophagy receptor that interacts with ABI3, ABI4, and ABI5 (Tarnowski *et al*., 2020). Expression pattern of *ATG7* (Figure S4) and *ABI5* (Albertos *et al*., 2015) co-localizes in the radicle zone, showing additional evidence of this relationship. These TFs have also been detected in ATG8 immunoprecipitation assays (Dagdas Y, personal communication). Moreover, in our co-IP assays, the ABI5–ATG8 interaction is observed *in vivo*, and it becomes more pronounced when the autophagy inhibitor (3-MA) is added to the medium. This indicates a direct link between ABI5 and autophagic processes in the early stages of development. We confirmed direct interactions between ATG8 and ABI3/ABI5 using Y2H assays (Fig. 9), supporting the possibility that these TFs are autophagy targets. Additionally, ABI5 degradation during germination follows a profile consistent with ATG8-mediated autophagy (Fig. 9). In *atg5* and *atg7*, this degradation is slower, aligning with their ABA hypersensitivity and delayed germination. Autophagy also affects bZIP67, a bZIP TF that corresponds to ABI5 homolog. In *atg7*, both bZIP67 levels and expression of its targets (*CRU1, CRU2, CRU3, AT2S1, FAE1*) are increased (Mendes *et al*., 2013; Fig. S10), suggesting autophagic regulation of bZIP67 during seed maturation. Altogether, our data support a model where autophagy modulates ABA responses during seed germination by controlling the abundance of key TFs, enabling rapid adaptation to environmental signals.

## Conclusions

The data presented here highlight the role of autophagy not only in the accumulation of reserves during seed maturation but also in the mobilization of storage compounds during early seed imbibition in *A. thaliana*. Our findings support the role of autophagy in cell vesicle trafficking in the radicle, contributing to the cellular dynamics of LD (lipid turnover) and PSV (protein storage and mobilisation). Our findings also indicate a role for autophagy in mediating the ABA response during early imbibition. This response involves the regulation of protein levels of selected TFs, such as ABI5 and bZIP67, which are pivotal in specific environmental responses. Consequently, autophagy emerges as a broad mechanism that controls how germinating seeds respond rapidly to environmental cues (Fig. 10). Therefore, autophagy likely plays a crucial role in the seed development, and germination programs and seed environmental perception (seed vigour) in *A. thaliana*. However, the question of whether these processes in *A. thaliana* can be extrapolated to other dicotyledonous species or even monocotyledonous species, with different seed anatomy and storage compounds, needs further investigation. Our work sheds light on seed autophagy and its fundamental importance in enhancing plant yield. Targeting autophagy genes associated with high seed yield and vigour will be crucial to achieving this goal.

**Fig. 10.**
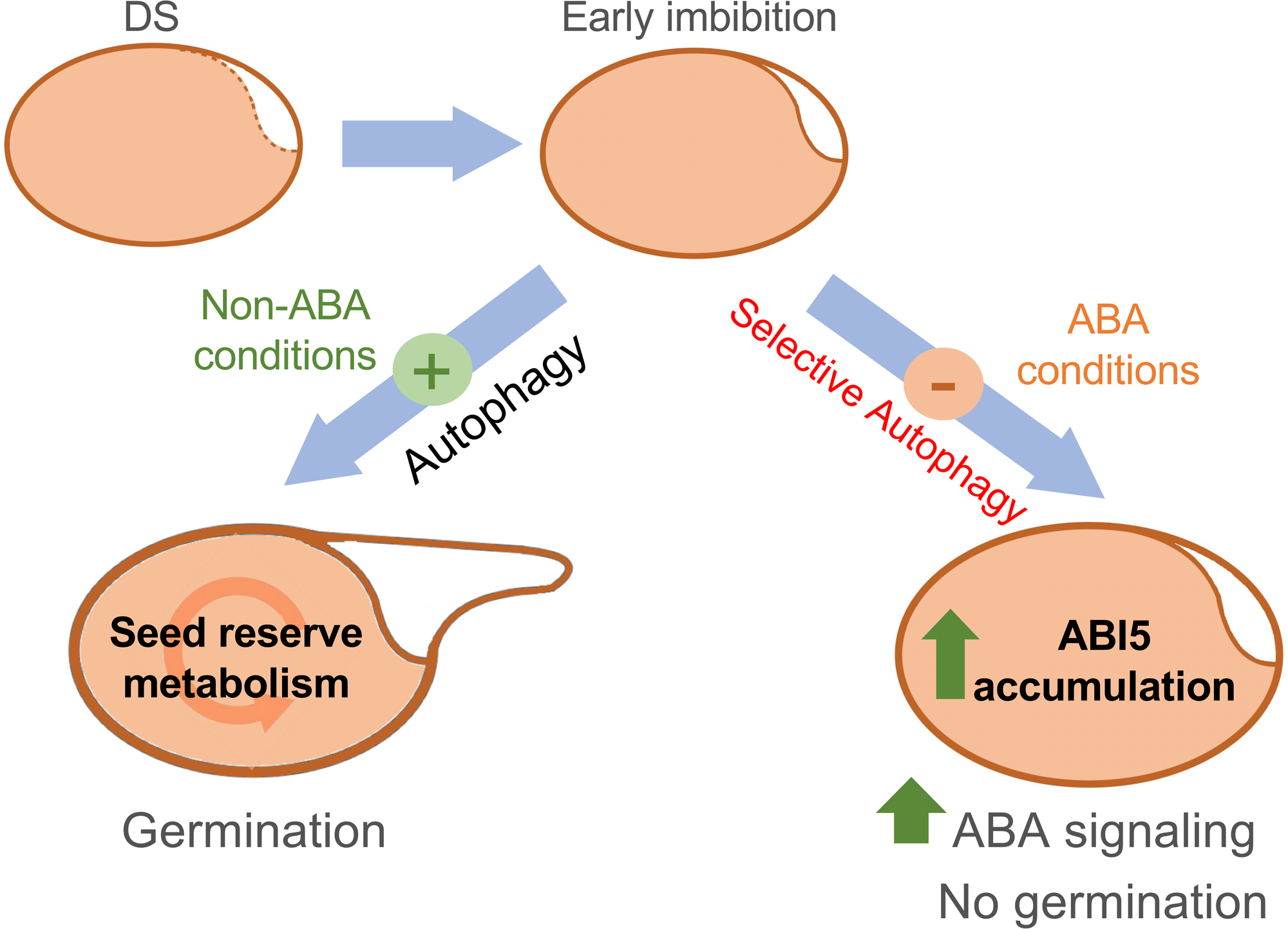
Proposed model of the integrative role of autophagy in early seed imbibition. According to our data, under non-stress conditions (non-ABA conditions), autophagy is crucial in controlling lipid turnover and protein storage mobilisation during germination. On the other hand, the ABA signal transduction pathway leads to modifications of downstream targets like ABI5, modulating their activity and participation in responses to environmental signals. The degradation of this kind of TF is critical for seed germination, and the contribution of a selective autophagy mechanism could contribute to modulate ABA signaling in response to adaptation to the environment.

## Acknowledgments

This work was supported by projects PID2023-148279OB-I00 and BIO2017-82873-R (PIs: R.I-F. and J.V.C and), Quinoa4Med (PRIMA-AEI, ref: PCI2022-132988; P.I.: R.I.F.), and PID2023-149447OB-I00, funded by MCIU/AEI/10.13039/ 501100011033, and project SA142P23 from the Regional Government of Castile and Leon and “Escalera de Excelencia” CLU-2018-04 co-funded by the FEDER Operative Program of Castile and Leon 2014–2020 Spain (to O.L.). Research grant funded by Fundación Memoria de D. Samuel Solórzano Barruso-Universidad de Salamanca Ref FS/8-2024 (IP: I. S-V.). We also thank the Spanish Research Network *EvoDevoSigNet* (RED2022-134917-T, MICIU).

The authors kindly thank Prof. A. Matilla and Rosina Saviaar (Mondego Science) for the critical reading of the manuscript.

## Author contributions

R.I.F. and J.V.C. conceived the project and reviewed the manuscript. R.I.F., J.V.C. and E.C. wrote the manuscript and prepared the figures with contributions of all authors. R.I.F. performed the bioinformatic analysis. E.C., M.A., and E.P. conducted all the immunohistochemical assays under the supervision of R.I.F. E.P. performed the expression analyses using qPCR. M.G.C. carried out the cloning of the *pATG5/7:uidA* lines and those used for the yeast two-hybrid assays under supervision of J.V.C. E.C. performed the Western blot assays for ABI5 and ABI3. O.L. and I.S.V. conducted the assays for bZIP67 and co-IPs. M.A.D. performed the germination assays.

## Data availability

The data obtained from the RNA sequencing (RNA-Seq) analysis have been uploaded to the Sequence Read Archive (SRA) of the National Center for Biotechnology Information (NCBI) under the reference PRJNA1190055. The codes used to perform the analysis with RStudio are deposited in a repository on GitHub (https://github.com/RaquelIglesias/Autophaghy-in-Arabidopsis-seed-imbibition).

## Legends to supporting figures, tables and data

**Table S1**. Sequences of primers used in this work.

**Data S1.** List of genes found in the AmiGO server using autophagy Gene Ontology term and their expression in siliques and seeds retrieved from the Gene Expression Tool at the Bio-Analytic Resource for Plant Biology (BAR) public database (https://bar.utoronto.ca/), expressed in GeneChip Operating System (GCOS) units (scale 0-10,000).

**Data S2.** RNA-seq differential expression analysis results (DESeq2) from the comparison of *atg7* vs Col-0 imbibed seeds at 24 hoi. Key columns include: Gene ID, Normalized counts (average normalized counts: BaseMean), log_2_FoldChange (expression change between conditions), p-value (statistical significance), adjusted p-value (padj; FDR corrected).

**Data S3. Gene Set Enrichment Analysis (GSEA) from the comparison of *atg7* vs Col-0 imbibed seeds at 24 hoi. (a-sheet):** Key columns include: Description Gene Set, NES (Normalized Enrichment Score), p-value (statistical significance), FDR q-value (false discovery rate), Core_enrichment (subset of genes driving the enrichment). **(b-sheet)** contains selected enriched gene sets and their corresponding results from the differential expression analysis: Lipid storage, seed maturation, and response to ER stress categories. Genes that belong to selected categories are listed and those meeting the cut-offs (*p*-adj < 0.05 and Log_2_FC ≥ |2.0|) are highlighted in green. Data represented in Fig. 4c as Lollipop plot. (**c-sheet**) shows the analysis of genes belonging to the Seed Maturation category in B-sheet using the ShinyGO platform (http://bioinformatics.sdstate.edu/go/) with the AGRIS Plant TF DB database, displaying the potential TFs that could regulate the expression of these genes. A column has been added in b-sheet marking the possible target genes of ABI5. (**d-sheet**) shows the expression analysis of genes belonging to the Lipid metabolism.

**Data S4. RNA-seq differential expression analysis results (DESeq2) from the comparison of atg7 vs Col-0 imbibed seeds at 96 hoi (1** μM ABA) and 48 hoi (DEGs responding to ABA specifically in the *atg7* mutant compared to Col-0, at comparable physiological states, see Fig. S5). Key columns include: Gene ID, Normalized counts (average normalized counts: BaseMean), log2FoldChange (expression change between conditions), p-value (statistical significance), adjusted p-value (padj; FDR corrected).

**Data S5. Gene Ontology and Transcription Factor Analysis in *atg7* vs. Col-0 in response to ABA upon seed germination.** Treatment. a-sheet: Gene ontology (GO) analysis using ShinyGO for genes significantly downregulated in Data S4 (*p*-adj ≤ 0.05 and Log_2_FC< - 1.0). The most significant gene categories are highlighted in red, particularly those related to ABA response and abiotic stress. **b-sheet**: Gene ontology (GO) analysis using ShinyGO for genes significantly upregulated in Data S4 (*p*-adj ≤ 0.05 and Log_2_FC> 1.0). As in sheet A, key gene categories are marked in red, focusing on ABA and abiotic stress-related categories. **c-sheet**: Transcription factor (TF) analysis using the AGRIS Plant TF database linked to ShyniGO platform for potential regulators of the downregulated genes of Sheet-A. Potential targets of ABI5 are highlighted in red. **d-sheet:** Transcription factor (TF) analysis for the upregulated genes in Data S4, using the same AGRIS database than in C-Sheet. Potential ABI5 target genes are marked in red. **e-sheet:** Gene ontology (GO) analysis of the potential ABI5 target genes obtained in C-Sheet, specifically focusing on genes categorized under “ABA response” (highlighted in red). F-sheet: Differentially Gene expression analysis results for the genes highlighted in red from sheets D and E (Data taken from Data S4). These results correspond to the Lollipop chart displayed in Fig. 8c, illustrating the expression levels of the selected genes.

