## Supplementary material for "The *autophagy-related genes AtATG5* and *AtATG7* influence reserve mobilisation and response to ABA during seed germination": Supplmentary Figures: Supp-Fig.pdf

### Figure S1

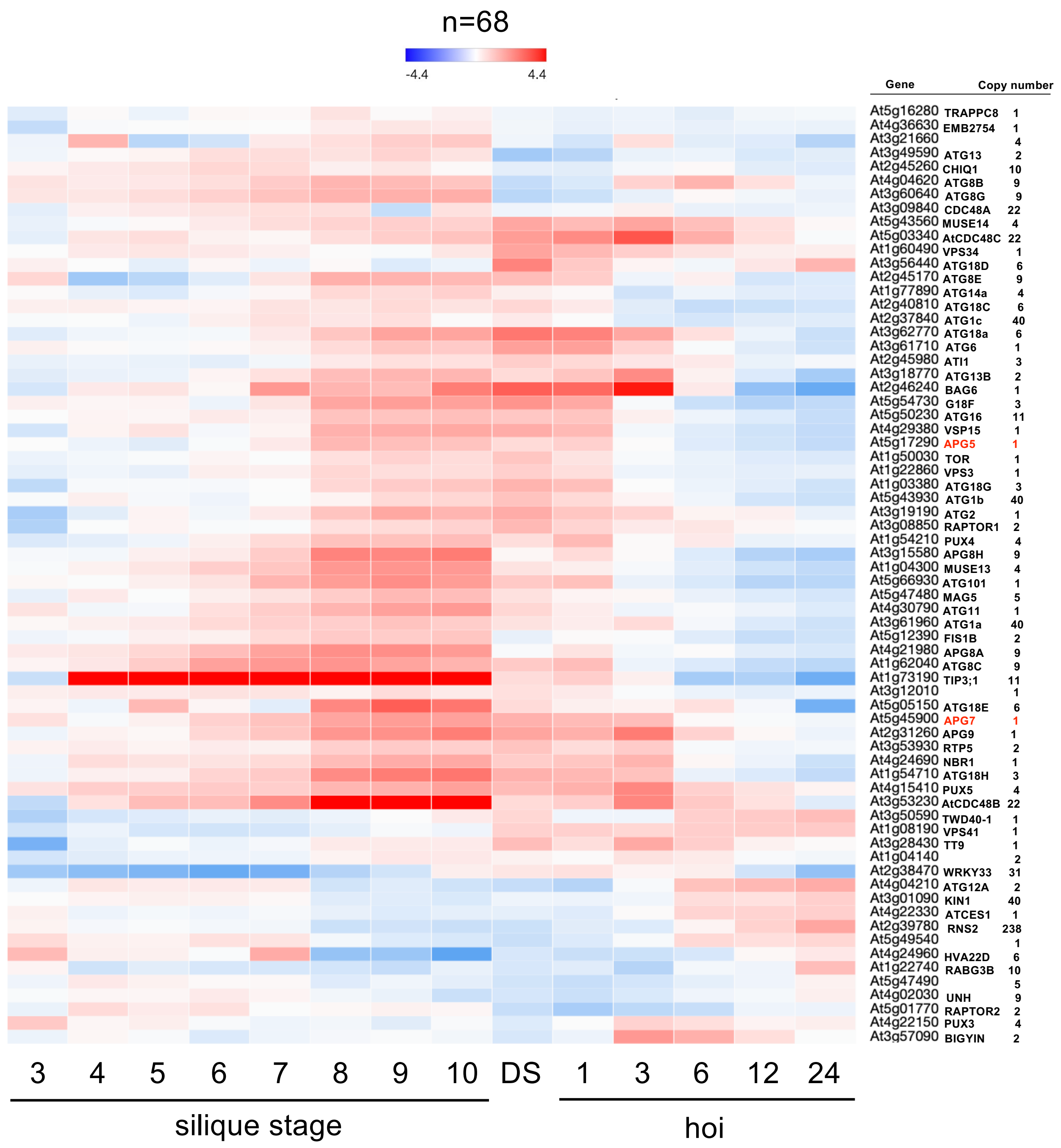
n=68
-4.4
4.4
Gene
Copy number
At5g16280 TRAPPC8 1
At4g36630 EMB2754 1
At3g21660 4
At3g49590 ATG13 2
At2g45260 CHIQ1 10
At4g04620 ATG8B 9
At3g60640 ATG8G 9
At3g09840 CDC48A 22
At5g43560 MUSE14 4
At5g03340 AtCDC48C 22
At1g60490 VPS34 1
At3g56440 ATG18D 6
At2g45170 ATG8E 9
At1g77890 ATG14a 4
At2g40810 ATG18C 6
At2g37840 ATG1c 40
At3g62770 ATG18a 6
At3g61710 ATG6 1
At2g45980 ATI1 3
At3g18770 ATG13B 2
At2g46240 BAG6 1
At5g54730 G18F 3
At5g50230 ATG16 11
At4g29380 VSP15 1
At5g17290 APG5 1
At1g50030 TOR 1
At1g22860 VPS3 1
At1g03380 ATG18G 3
At5g43930 ATG1b 40
At3g19190 ATG2 1
At3g08850 RAPTOR1 2
At1g54210 PUX4 4
At3g15580 APG8H 9
At1g04300 MUSE13 4
At5g66930 ATG101 1
At5g47480 MAG5 5
At4g30790 ATG11 1
At3g61960 ATG1a 40
At5g12390 FIS1B 2
At4g21980 APG8A 9
At1g62040 ATG8C 9
At1g73190 TIP3;1 11
At3g12010 1
At5g05150 ATG18E 6
At5g45900 APG7 1
At2g31260 APG9 1
At3g53930 RTP5 2
At4g24690 NBR1 1
At1g54710 ATG18H 3
At4g15410 PUX5 4
At3g53230 AtCDC48B 22
At3g50590 TWD40-1 1
At1g08190 VPS41 1
At3g28430 TT9 1
At1g04140 2
At2g38470 WRKY33 31
At4g04210 ATG12A 2
At3g01090 KIN1 40
At4g22330 ATCES1 1
At2g39780 RNS2 238
At5g49540 1
At4g24960 HVA22D 6
At1g22740 RABG3B 10
At5g47490 5
At4g02030 UNH 9
At5g01770 RAPTOR2 2
At4g22150 PUX3 4
At3g57090 BIGYIN 2
3 4 5 6 7 8 9 10 DS 1 3 6 12 24
silique stage
hoi

### Figure S2

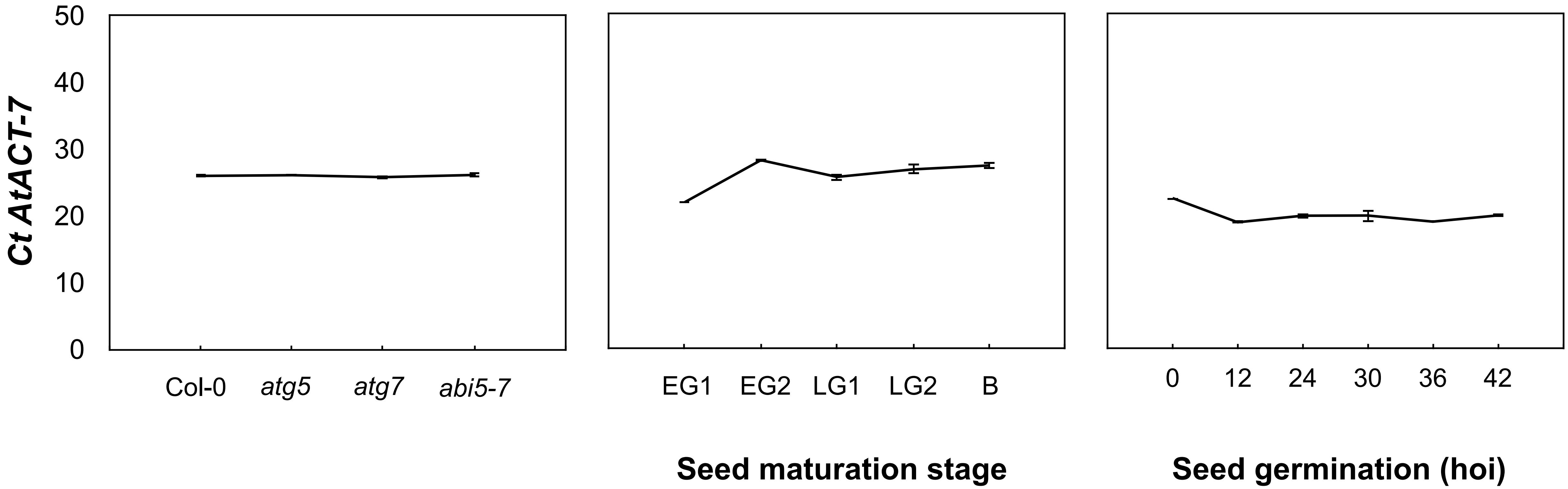

Ct AtACT-7
50
40
30
20
10
0
Col-0
atg5
atg7
abi5-7
EG1
EG2
LG1
LG2
B
Seed maturation stage
0
12
24
30
36
42
Seed germination (hoi)

### Figure S3

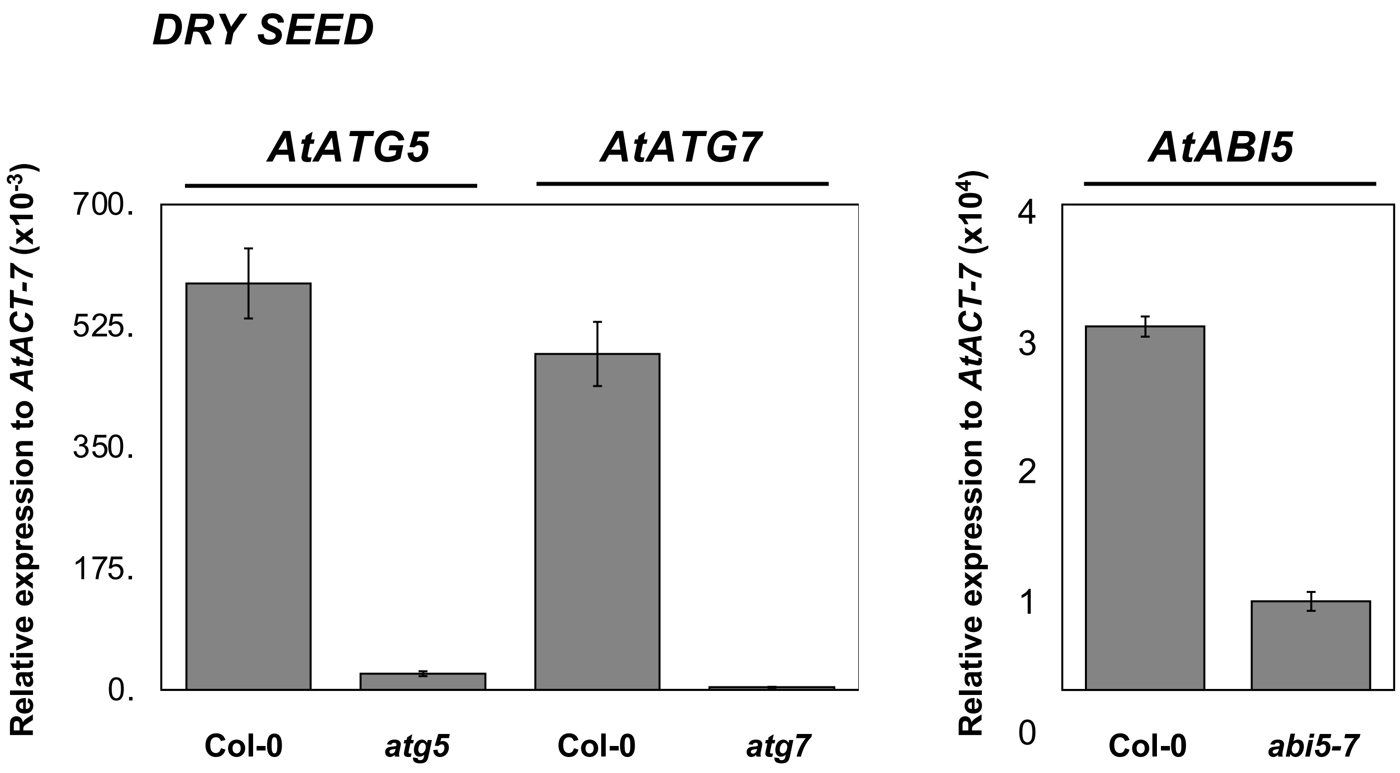

DRY SEED
AtATG5
AtATG7
AtABI5
Relative expression to AtACT-7 (x10-3)
700.
525.
350.
175.
0.
Col-0
atg5
Col-0
atg7
Relative expression to AtACT-7 (x104)
4
3
2
1
0
Col-0
abi5-7

### Figure S4

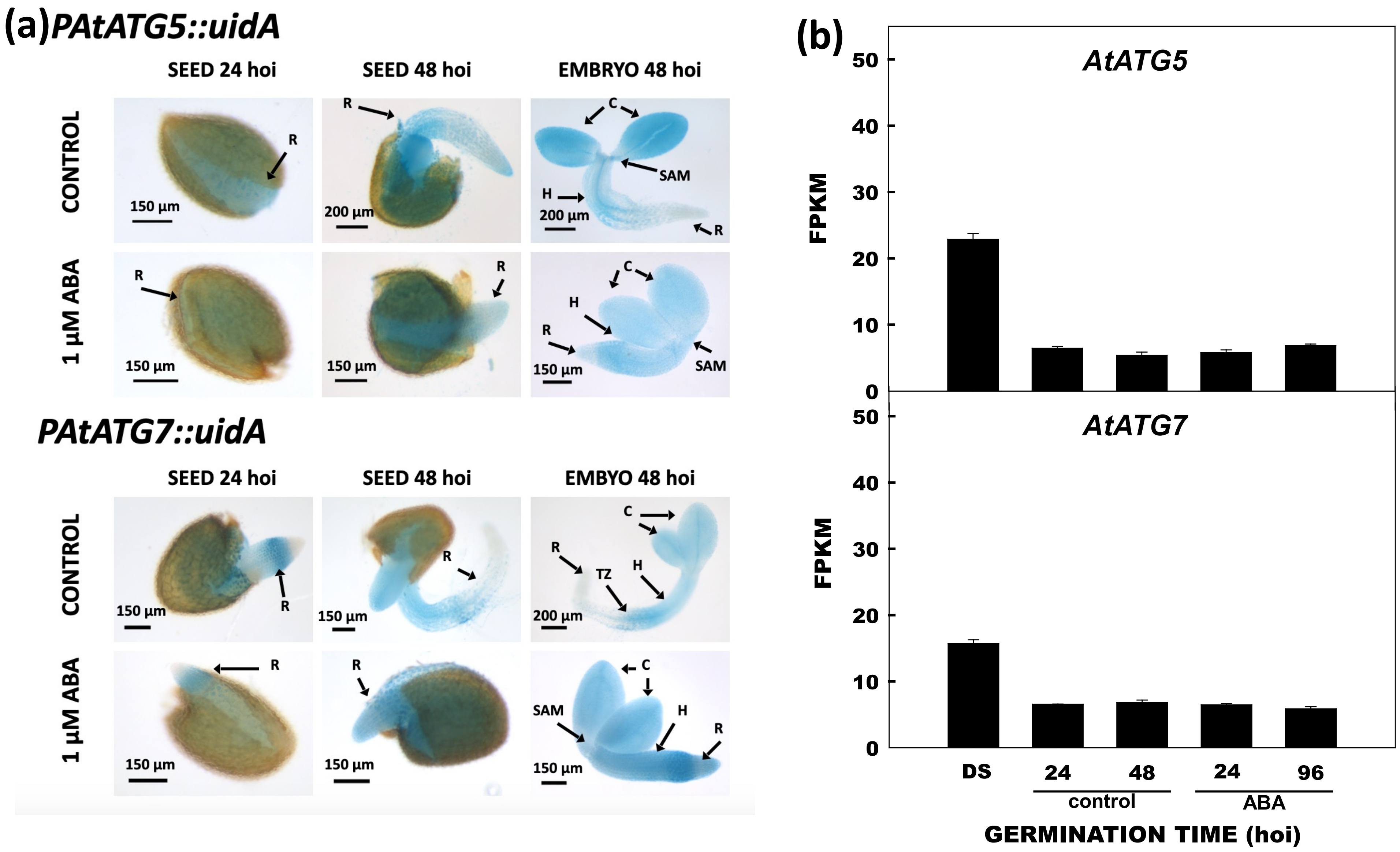

(a)PAtATG5::uidA
SEED 24 hoi
SEED 48 hoi
EMBRYO 48 hoi
CONTROL
1 µM ABA
R
C
SAM
H
150 µm
200 µm
PAtATG7::uidA
SEED 24 hoi
SEED 48 hoi
EMBYO 48 hoi
CONTROL
1 µM ABA
TZ
150 µm
200 µm
(b)
AtATG5
AtATG7
FPKM
0
10
20
30
40
50
DS
24
48
24
96
control
ABA
GERMINATION TIME (hoi)

### Figure S5

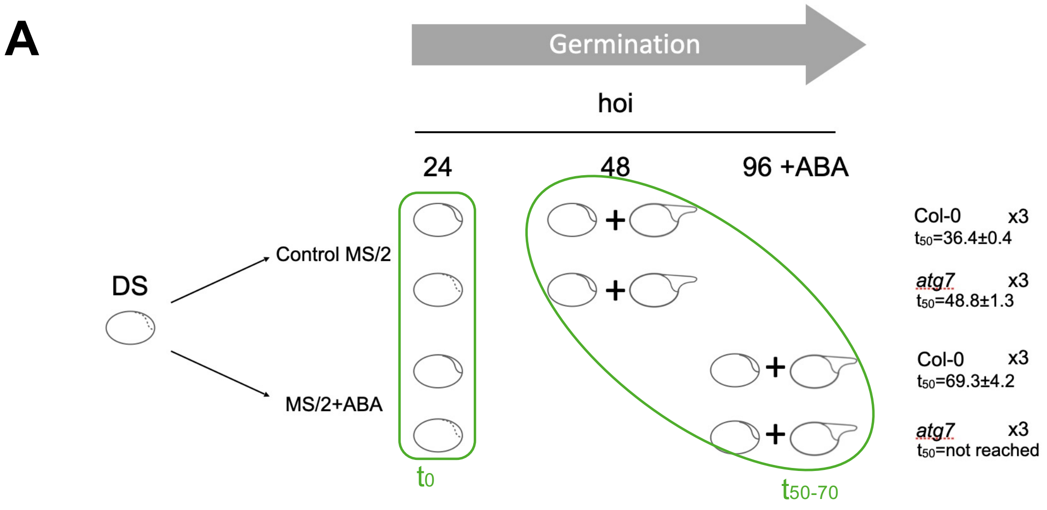
A
Germination
hoi
24
48
96 +ABA
DS
Control MS/2
MS/2+ABA
Col-0 x3
t50=36.4±0.4
atg7 x3
t50=48.8±1.3
Col-0 x3
t50=69.3±4.2
atg7 x3
t50=not reached
t0
t50-70

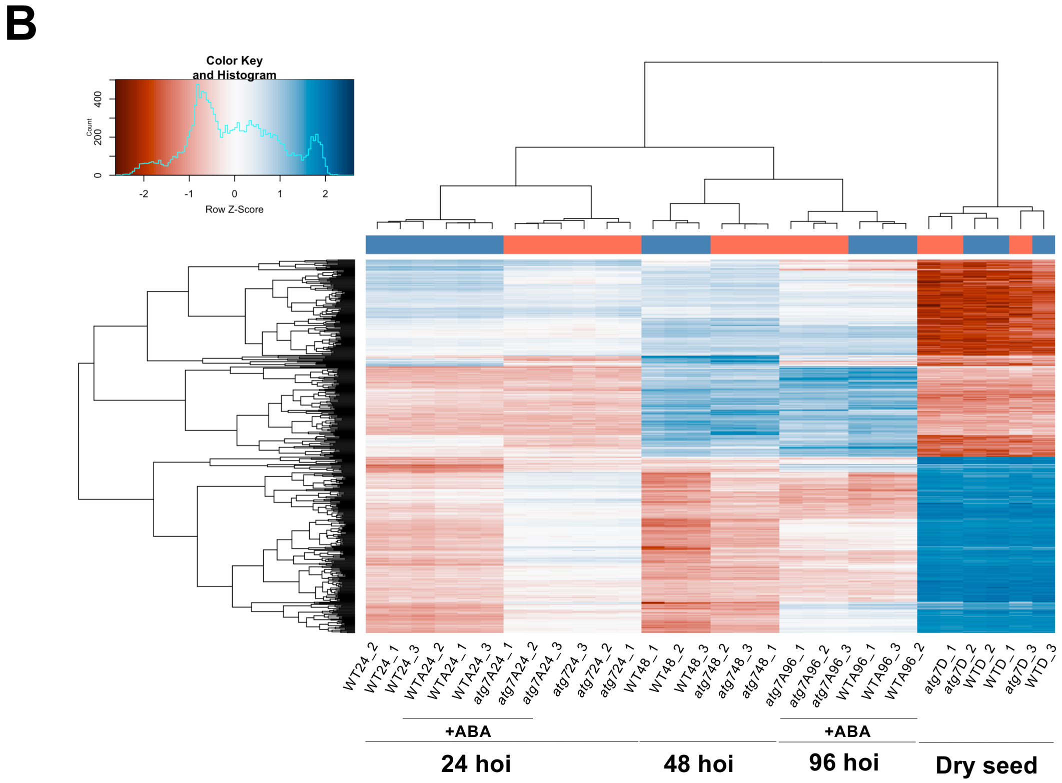
B
Color Key and Histogram
Count
Row Z-Score
WT24_2
WT24_1
WT24_3
WTA24_2
WTA24_1
WTA24_3
atg7A24_1
atg7A24_2
atg7A24_3
atg724_3
atg724_2
atg724_1
WT48_1
WT48_2
WT48_3
atg748_2
atg748_3
atg748_1
atg7A96_1
atg7A96_2
atg7A96_3
WTA96_1
WTA96_3
WTA96_2
atg7D_1
atg7D_2
WTD_2
WTD_1
atg7D_3
WTD_3
+ABA
+ABA
24 hoi
48 hoi
96 hoi
Dry seed

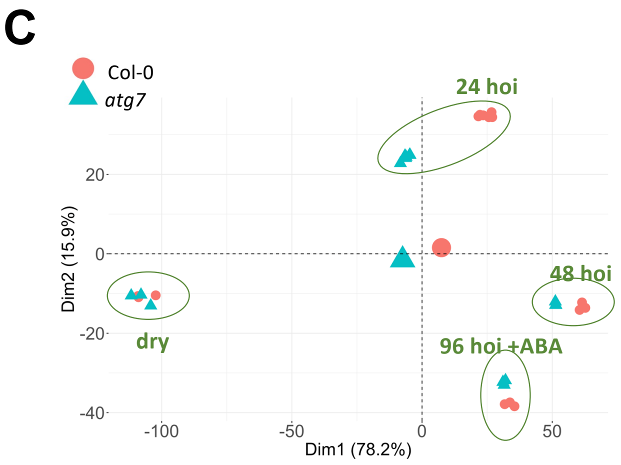
C
Col-0
atg7
24 hoi
48 hoi
dry
96 hoi +ABA
Dim2 (15.9%)
Dim1 (78.2%)

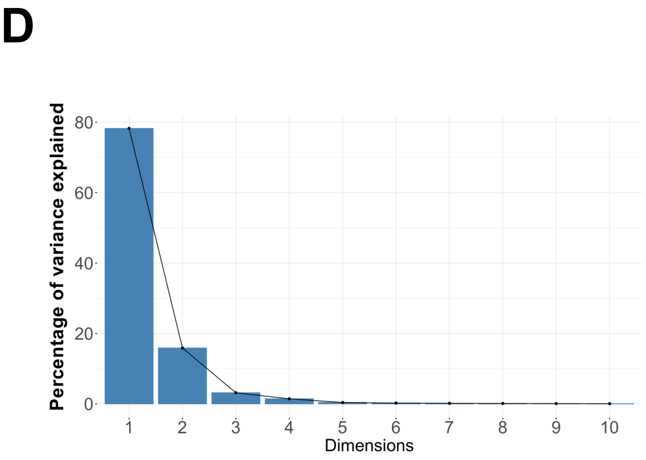
D
Percentage of variance explained
Dimensions

### Figure S6

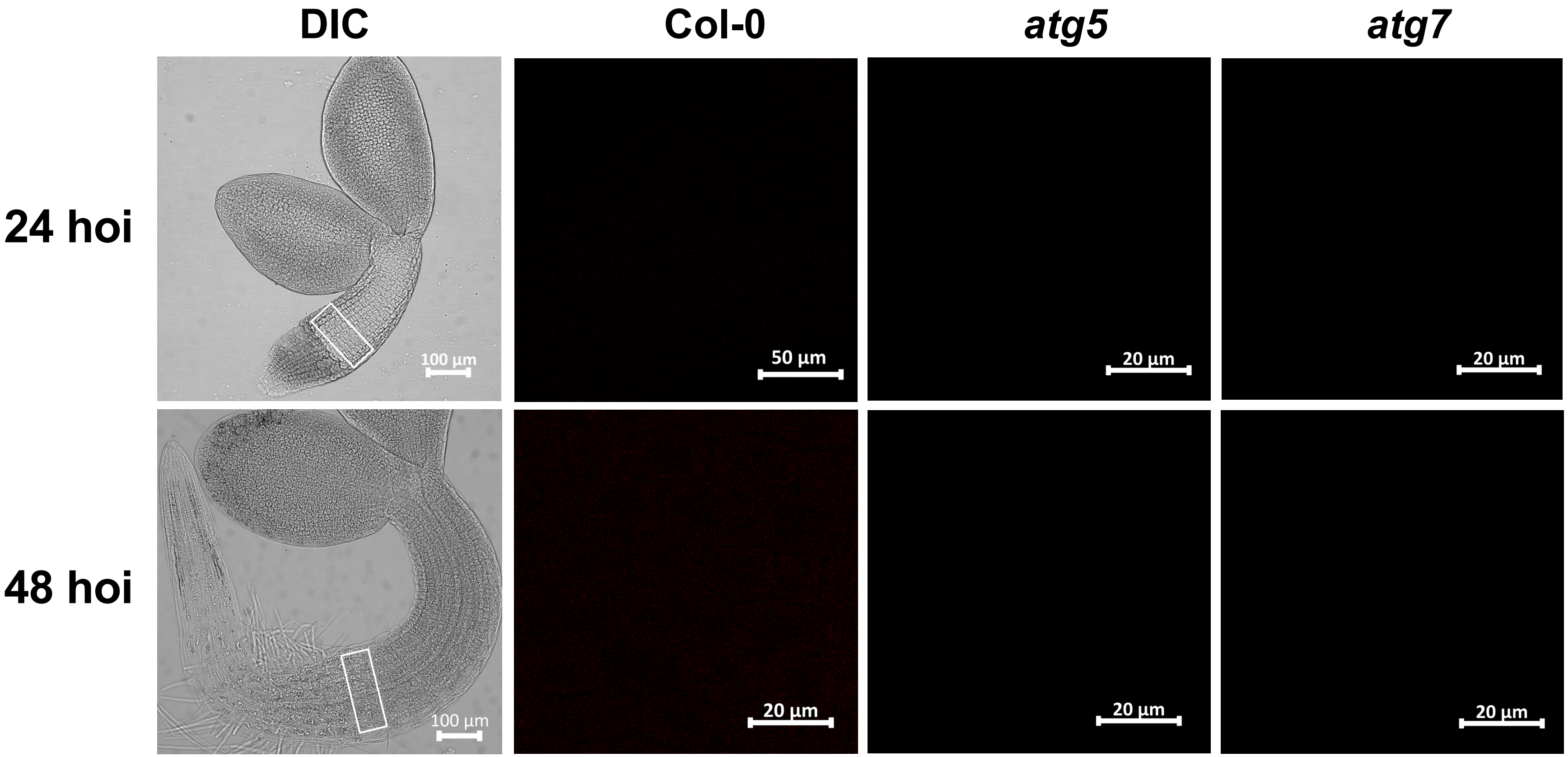

DIC
Col-0
atg5
atg7
24 hoi
48 hoi
100 µm
50 µm
20 µm
20 µm
100 µm
20 µm
20 µm
20 µm

### Figure S7

(a)

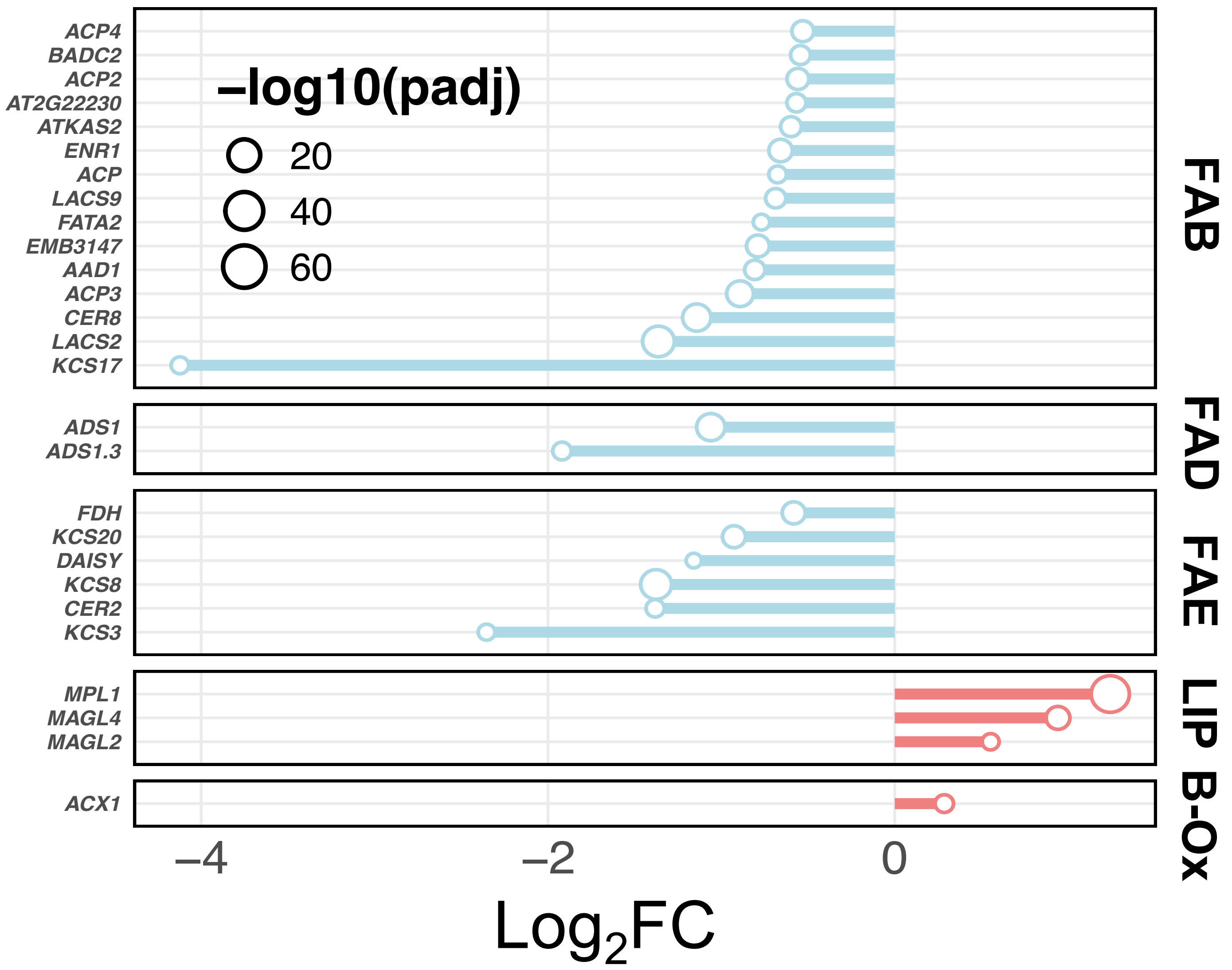


(c)

| Gene name | Enzymatic Activity |
|---|---|
| *ACX* | EC:1.3.3.6 |
| *AT2G22230* | EC:4.2.1.59 |
| *ATKAS2* | EC 2.3.1.179 |
| *CER8* | EC:6.2.1.3 |
| *DAISY* | EC:2.3.1.199 |
| *EMB3147* | EC: 2.3.1.39 |
| *ENR1* | EC:1.3.1.9 and EC:1.3.1.10 |
| *FATA2* | EC:3.1.2.14 |
| *FDH* | EC:2.3.1.199 |
| *KCS20* | EC:2.3.1.199 |
| *KCS3* | EC:2.3.1.199 |
| *KCS8* | EC:2.3.1.199 |
| *LACS2* | EC:6.2.1.3 |
| *LACS9* | EC:6.2.1.3 |
| *MAGL2* | EC:3.1.1.23 |
| *MAGL4* | EC:3.1.1.23 |
| *MPL1* | EC:3.1.1.3 |

(b)

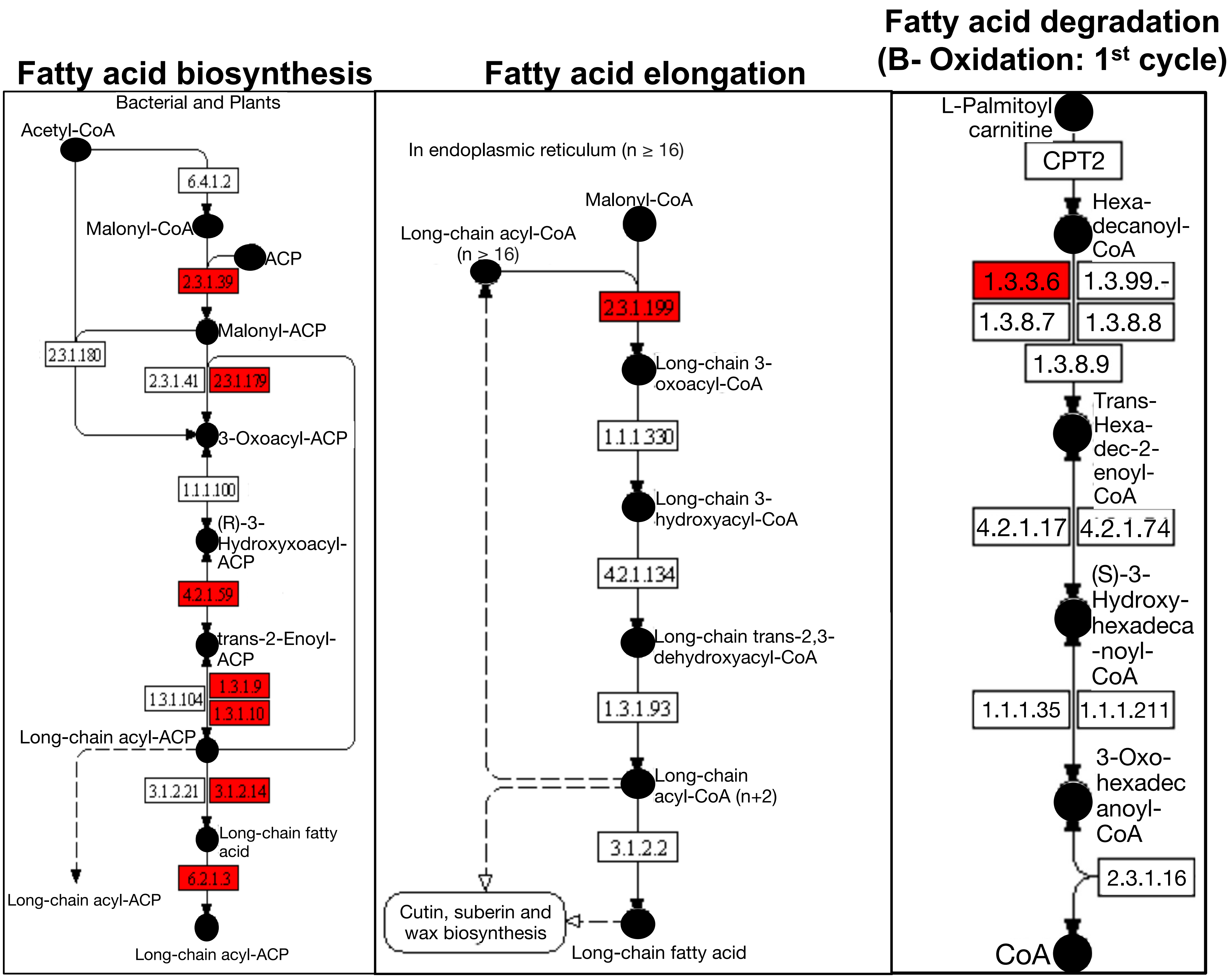

Figure S8

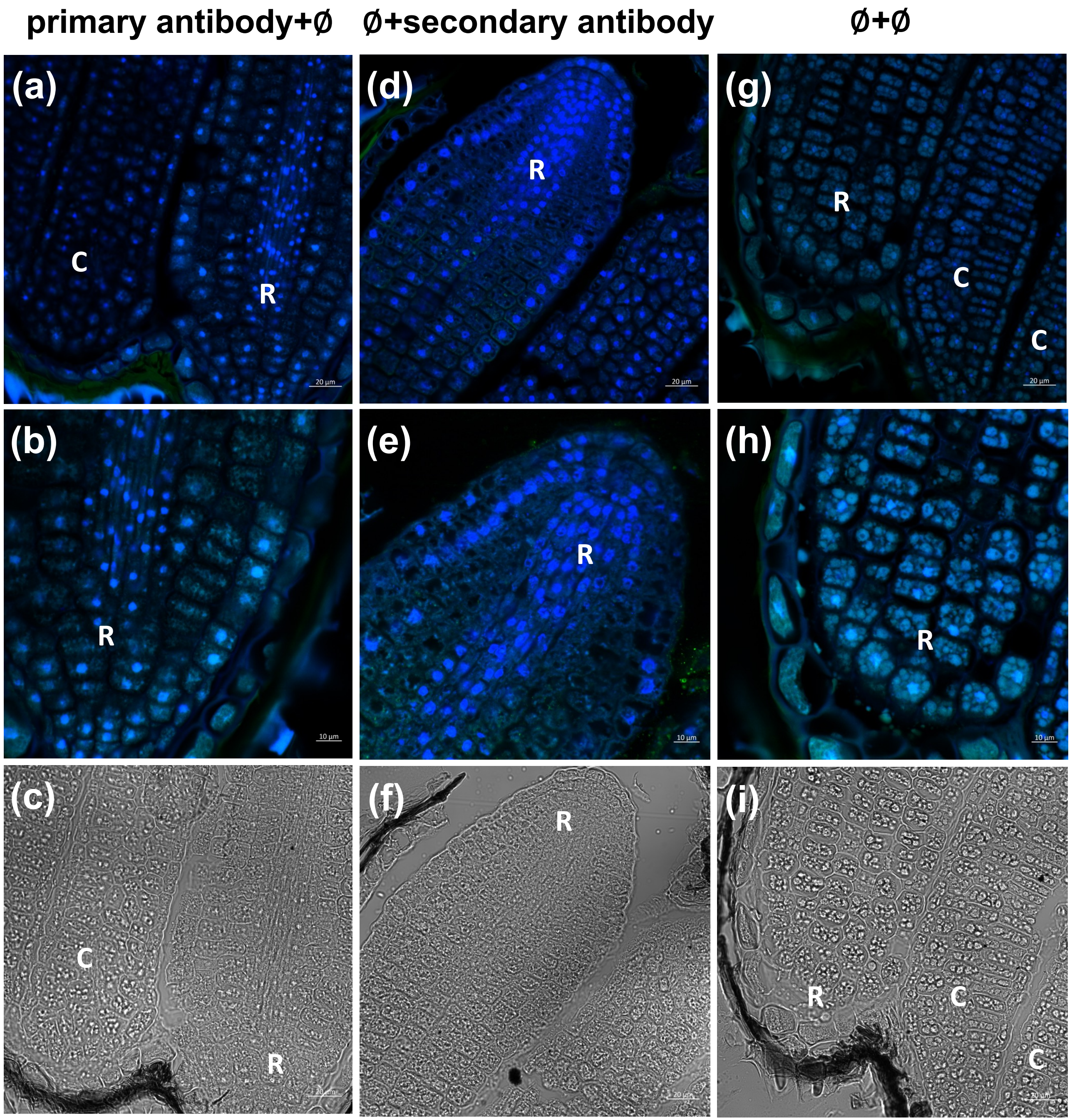
primary antibody+Ø
Ø+secondary antibody
Ø+Ø
(a)
C
R
(b)
R
(c)
C
R
(d)
R
(e)
R
(f)
R
(g)
R
C
C
(h)
R
(i)
R
C
C

### Figure S9

(a)

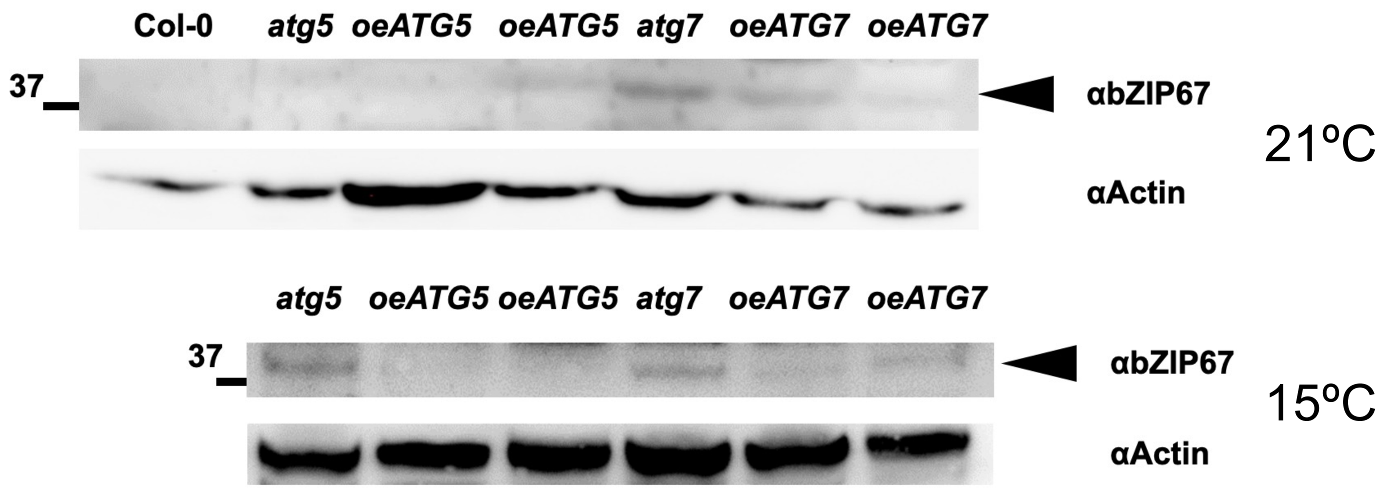


(b)

MSVFESETSNFHVYNNHEIQTQPQMQTFLSEEEPVGRQNSI
LSLTLDEIQMKSGKSFGAMNMDEFLANLWTTVEENDNEGGG
AHNDGEKPAVLPRQGSLSLPVPLCKKTVDEVWLEIQNGVQQ
HPPSSNSGQNSAENIRRQQTLGEITLEDFLVKAGVVQEPLK
TTMRMSSSDFGYNPEFGVGLHCQNQNNYGDNRSVYSENRPF
YSVLGESSSCMTGNGRSNQYLTGLDAFRIKKRIIDGPPEIL
MERRQRRMIKNRESAARSRARRQAYTVELELELNNLTEENT
KLKEIVEENEKKRRQEIISRSKQVTKEKSGDKLRKIRRMAS
AGW

(c)

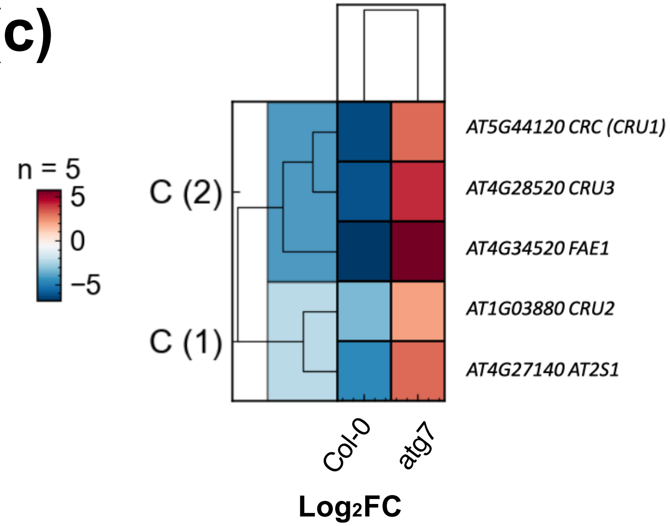

### Figure S10

#### ABI3

MKSLHVAANAGDLAEDCGILGGDADDTVLMDGIDEVGREIWLDDHGGDNNHVHGHQDDDL
IVHHDPSIFYGDLPTLPDFPCMSSSSSSSTSPAPVNAIVSSASSSSAASSSTSSAASWAI
LRSDGEDPTPNQNQYASGNCDDSSGALQSTASMEIPLDSSQGFGCGEGGGDCIDMMETFG
YMDLLDSNEFFDTSAIFSQDDDTQNPNLMDQTLERQEDQVVVPMMENNSGGDMQMMNSSL
EQDDDLAAVFLEWLKNNKETVSAEDLRKVKIKKATIESAARRLGGGKEAMKQLLKLILEW
VQTNHLQRRRTTTTTTNLSYQQSFQQDPFQNPNPNNNNLIPPSDQTCFSPSTWVPPPPQQ
QAFVSDPGFGYMPAPNYPPQPEFLPLLESPPSWPPPPQSGPMPHQQFPMPPTSQYNQFGD
PTGFNGYNMNPYQYPYVPAGQMRDQRLLRLCSSATKEARKKRMARQRRFLSHHHRHNNNN
NNNNNNQQNQTQIGETCAAVAPQLNPVATTATGGTWMYWPNVPAVPPQLPPVMETQLPTM
DRAGSASAMPRQQVVPDRRQGWKPEKNLRFLLQKVLKQSDVGNLGRIVLPKKEAETHLPE
LEARDGISLAMEDIGTSRVWNMRYRFWPNNKSRMYLLENTGDFVKTNGLQEGDFIVIYSD
VKCGKYLIRGVKVRQPSGQKPEAPPSSAATKRQNKSQRNINNNSPSANVVVASPTSQTVK

#### ABI5

MVTRETKLTSEREVESSMAQARHNGGGGGENHPFTSLGRQSSIYSLTLDEFQHALCENGKN
FGSMNMDEFLVSIWNAEENNNNQQQAAAAAGSHSVPANHNGFNNNNNNGGEGGVGVFSGGS
RGNEDANNKRGIANESSLPRQGSLTLPAPLCRKTVDEVWSEIHRGGGSGNGGDSNGRSSSS
NGQNNAQNGGETAARQPTFGEMTLEDFLVKAGVVREHPTNPKPNPNPNQNQNPSSVIPAAA
QQQLYGVFQGTGDPSFPGQAMGVGDPSGYAKRTGGGGYQQAPPVQAGVCYGGGVGFGAGGQ
QMGMVGPLSPVSSDGLGHGQVDNIGGQYGVDMGGLRGRKRVVDGPVEKVVERRQRRMIKNR
ESAARSRARKQAYTVELEAELNQLKEENAQLKHALAELERKRKQQVKTPIEFALLRWLQYF
GQKKTTKWNCWFMVAVF
